# Transcription Factor Target Gene Network governs the Logical Abstraction Analysis of the Synthetic Circuit in Leishmaniasis

**DOI:** 10.1101/151779

**Authors:** Milsee Mol, Dipali Kosey, Ramanamurthy Bopanna, Shailza Singh

## Abstract

Stochastic variations in the transcription factor target gene network influences the dynamics of protein levels. The mathematical model built, here, is useful to study the cytokine response and the resulting dynamical patterns in leishmaniasis. The fluctuations produced affect the expression rate of its target in the regulatory synthetic circuit designed by means of a negative feedback loop insertion. Thus, the faster the response time, smaller is the fluctuation of the system observed and it can be justified that the TFTG network embedded can be understood with the recurring dynamics of the designed synthetic circuit.

**List of Abbreviations**
TF: transcription factor
PPARα: peroxisome proliferator-activated receptor-
FAs: fatty acids
DCs: dendritic cells
TFTG network: Transcription Factor Target Gene Network
IFNg: interferon g
VL: visceral leishmaniasis
CL: cutaneous leishmaniasis
G-MCF: granulocyte macrophage colony-stimulating factor
IL: interleukin
PKC: Protein Kinase C
PBC: Periodic boundary condition
NPT: Normal Pressure temperature
NVT: Normal Volume Temperature
MD: Molecular Dynamics
RMSD: root mean square deviation
RMSF: root mean square fluctuation
LB: Luria Berttini
I: Infection
CT: Chimeric PKC
CTI: Chimeric PKC + Infection
CTM: Chimeric PKC + Miltefosine
CTIM: Chimeric PKC + Infection + Miltefosine
CC: Closeness centrality
IPTG: Isopropyl β-D-1-thiogalactopyranoside

## 1. Introduction

Leishmaniasis is a group of complex disease caused by a protozoan parasite belonging to the genus *Leishmania*. There are almost 20 *Leishmania* species that can infect humans by the bite of infected female phlebotomine sand-fly belonging to the genus *Lutzomiya* or *Phlebotomus*. It is restricted to the tropical areas of the world and over 98 countries are endemic for leishmaniasis. It is estimated that approximately 0.2 to 0.4 million cases of visceral leishmaniasis (VL) and 0.7 to 1.2 million cases of cutaneous leishmaniasis (CL) occur each year worldwide (www.who.in). *Leishmania* shows a digenetic life cycle alternating between the vector sandfly and mammalian host (www.cdc.gov). Treatment options in leishmaniasis are few due to limited understanding of the biology of *Leishmania* spp. and the host parasite interaction. The available chemotherapeutics like pentamonials, amphotericin B, pentamidine and miltefosine cannot be considered ideal due to their side effects such as high toxicity, extended duration of treatment and economic feasibility. More than often the parenteral mode of drug administration leads to treatment abandonment ^1–8^.

Since, *Leishmania* establish an infection in their host macrophages if they are capable of evading and supressing the adaptive immune response, immune-stimulation could be another approach to treat leishmaniasis. Immuno modulators like endogenous interferon g (IFNg), granulocyte macrophage colony-stimulating factor (G-MCF), interleukin (IL) 12, bacterial and fungal derivatives like muramyl dipeptide, trehalose dimycolate, and glucan, synthetic compounds like levamisole, lipoidal amine, cimetidine, tuftsin, polyinosinic-polycytidylic acid, imiquimod and tucaresol have shown promise in the treatment of leishmaniasis ^9^. Liposomes and nanoparticles have also been developed as alternative treatment options for leishmaniasis. Liposomal formulation of amphotericin B, Ambisome, stearylamine liposomal formulation containing sodium stibogluconate, oleylphosphocholine (OlPC) encapsulated in liposomes have shown some leishmanicidal action. Similarly, nanoparticles of Amphotericin B attached to functionalized carbon nanotubes or chitosan-chondroitin sulphate nanoparticles have also shown leishmanicidal activity ^10–12^.

Since immunobiology of the parasite is not well understood, vaccine development is also a major challenge.

### Immunobiology of Leishmaniasis

Leishmaniasis is termed as an immunologically highly complex disease. The disease may be cured or exacerbated depending on genetic variations of the mammalian host and *Leishmania* species, inoculum size and bite size received. Upon entry into the mammalian host, the leishmanial promastigotes are engulfed by the dendritic cells (DCs), neutrophils and macrophages. The infected macrophages may produce IL12 which activates the natural killer (NKs) cells, which in turn produce IFNg followed by Th1 activation. Tumour necrosis factor-a (TNFa) synergizes with IFNg to activate infected macrophages that kill the parasites primarily through oxidative mechanisms. While on the other hand the parasite is also capable of initiating Th2 response for disease progression by modulating the immune signalling of the macrophages. Th2 response is characterized by the early production of IL4, IL10 and IL13, absence of synthesis of IL12. Thus, in leishmaniasis, macrophages play a dual role; they represent an important cell population, i.e. M1 responsible for killing of the parasites by activating the Th1 adaptive immune response. Moreover, the major site of parasite replication in which Th2 adaptive immune response is dominant is activated by the M2 macrophages ^13–20^ (**Figure 1**).

**Figure 1:**
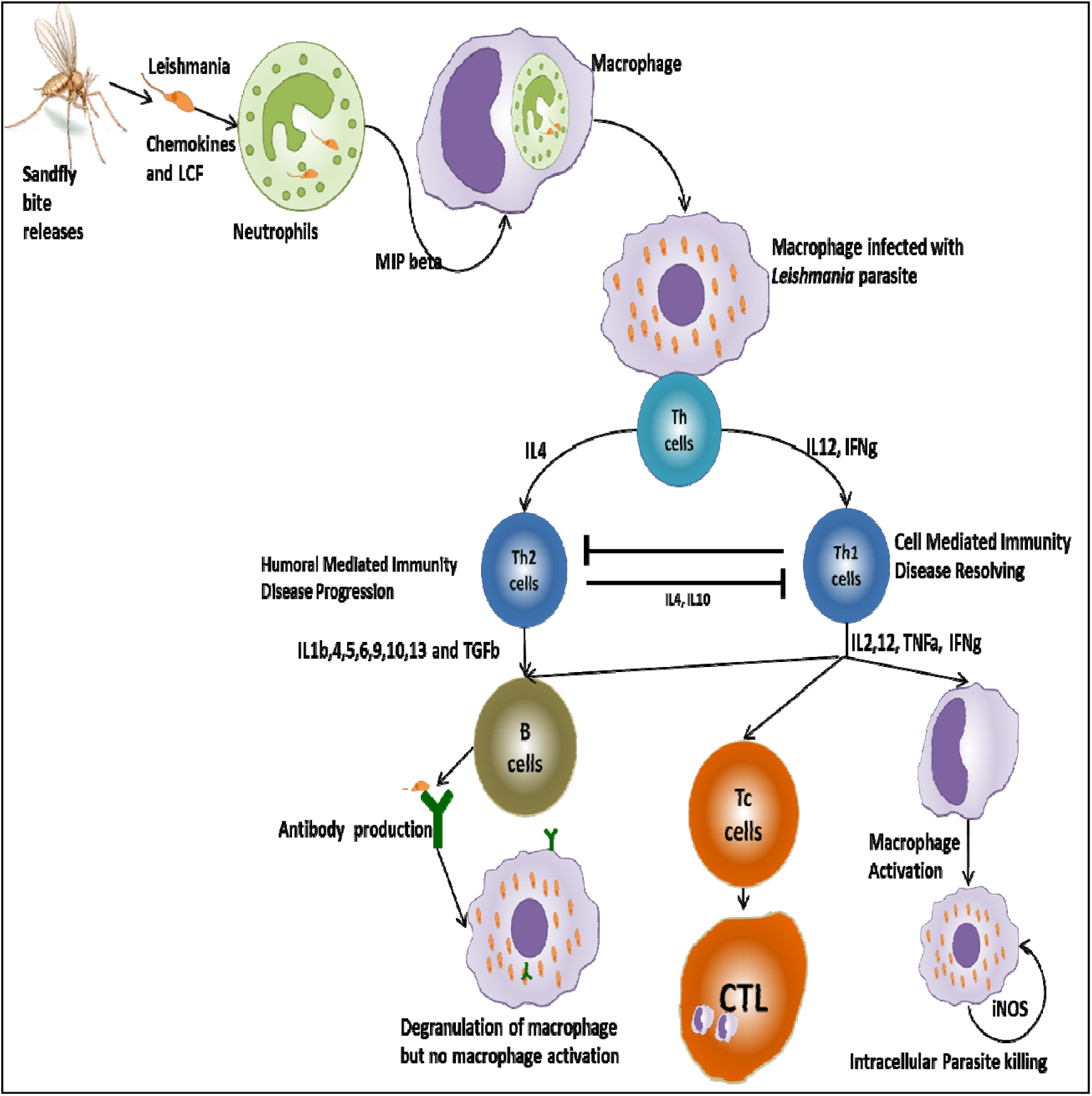
*Leishmania* infection and its Th1-Th2 tweaking via macrophages.

### Macrophage Plasticity

The macrophages are flexible and continually changeable, and they can change from one functional phenotype to another in response to change in cytokine stimulation. This ability of macrophages has been tapped for application in therapeutics against different pathophysiological conditions like helminth infection ^21^ or muscle regeneration ^22,23^. Small pharmaceutical molecules are used to reverse the stressed cellular output from a diseased state to a healthy state, by acting on the intermediates of cellular signaling pathways, for example, small molecule inhibitors or monoclonal antibodies against receptor tyrosine kinases (RTKs), or other proteins with catalytic activity as applied in cancer ^24,25^. Some small molecules can change the anti-inflammatory phenotypic state to a pro-inflammatory phenotypic state and vice versa, for example, VX-765, a caspase1 inhibitors blocks the processing and release of the active forms IL1beta and IL18, showing a potent anti-inflammatory activity and has been applied in conditions such as psoriasis and Muckle-Wells syndrome ^26^. Active Salt-inducible kinases (SIKs), results in diminished IL10 production by phosphorylation of CRTC3 (CREB-regulated transcriptional co-activator 3). Small molecules like dasatnib and bosutinib, suppress the SIK activity by increasing the cAMP levels and potentiates IL10 production by macrophages and DCs. These molecules are being tested in condition of inflammatory bowel disease (IBD) ^27^. Autoimmune disorders like rheumatoid arthritis, IBD, psoriasis and septic shock syndrome can be treated by blocking the MAPKs at multiple levels in the signal transduction, resulting in reduced pro-inflammatory cytokine signaling ^28^.

As with small molecules changing the signal transduction for a healthy cell transition, signaling pathways can be rewired using engineered proteins embedded into a synthetic circuit. In particular, synthetic circuits capable of integrating multiple input signals, for example, AND, NAND and NOR logic gates, that can recognize complex cellular environment like disease-specific biomarkers, enhancing the specificity and accuracy of biological control.

Though synthetic biology is in its infancy, many proof of concept design principles are being created and tested *in vitro* and *in vivo*, inspired from systems biology studies. A binary synthetic transcription factor (TF) with fused domains from peroxisome proliferator-activated receptor-α (PPARα) and bacterial DNA-binding repressor TtgR was constructed for the treatment of diet induced obesity in mice. PPARα, in the presence of fatty acids (FAs) recruits’ co-activators and in the absence of fatty acids recruits co-repressors. Together with PPARα and FAs, the TtgR driven promoter expresses an appetite suppressing peptide hormone pramlintide. These encapsulated engineered cells when implanted in mice with diet-induced obesity, showed significant reduction in food consumption and blood lipid levels^29^. Circuits that can be controlled using light waves for therapeutic purposes are another appealing avenue. In an optogenetic device, shGLP-1 hormone with potent glucose homeostasis-modulating characteristics was put under transcriptional control of melanopsin, a blue light sensor protein, which transforms a light energy to change in membrane potential (due to Ca^2+^ influx) triggering an intracellular signaling cascade. This cascade activates a TFS via PKC controlling the expression of the hormone. The circuit was encapsulated cultured human cells and implanted in a human type II diabetes mouse model that improved glucose homeostasis ^30^.

Uric acid homeostasis was achieved by using a prosthetic gene network to deal with gout. In this gene network, urate oxidase is under direct control of bacterial uric acid sensor HucR. In the presence of uric acid, the HucR sensor leaves its target DNA motif resulting in the expression of urate oxidase. Thus, the synthetic circuit acts as a sensor and stabilizes the blood urate concentration. Also it has been shown that, the urate sensor shows a homeostatic action in urate oxidase-deficient mice ^31^.

Infectious disease immuno-biology is a complex phenomenon and needs attention in terms of designing appropriate methods and tools for combat. Systems and synthetic biology approaches can be applied to such systems and this paper has tried to address the application of synthetic biology for modulating the immune response towards the intracellular parasite *L. major* causing CL.

Systems approach to study the signaling cascade in macrophages, modulated by the *Leishmania* parasite needs emphasis with respect to its role in modulating the host signaling mechanism. When a macrophage is infected with *Leishmania,* multiple modulating signal inputs arrive simultaneously and/or sequentially, subsequently integrated for an anti-inflammatory pathophysiology in leishmaniasis. Novel biological devices can be constructed, that revert these signals for a pro inflammatory leishmanicidal response to curb infection.

In our earlier study the CD14, TNF and EGFR pathways are considered with in a systems purview for reconstruction. Network metrices indicate that the reconstructed signaling cascade is a real world network with common crosstalk points like the MKK1/2, MKK3/6, MKK4/7 and IKK–NFkB, which are central to modulations for different phenotypic response ^32^. These crosstalk points play an important role in leishmaniasis which is evident from large body of published literature. Some chemotherapeutics agents like miltefosine and some plant extracts act via modulation of some of these crosstalk mediators. But increasing incidence of drug resistance opens avenues for synthetic circuits to apply for immune modulation using these crosstalk points. For successful application of the synthetic circuits, it is pertinent to know the gene expression pattern of the signaling pathway which can be self-modulated. Henceforth, the TFTG network was reconstructed for the corresponding TFs that could be activated via the CD14-TNF-EGFR pathway, which is discussed subsequently.

## 2. Materials and Methods

### Reconstruction of TFTG network and Node selection

Since there are no direct evidence for the transcription factor and their corresponding target genes in leishmaniasis, knowledge from various databases and literature were collected and a graph based analysis was implemented on the reconstructed TFTG pair.

There were nine TFs that could be related to *Leishmania* infection that are part of the reconstructed CD14-TNF-EGFR signaling network namely Creb1, Elk1, Ets1, Fos, Jun, Myc, Nfkb, Stat1 and Stat3. The TFTG network reconstructed consists of possibly all the genes that are involved in an immune response in leishmaniasis. The initial listing of transcription factor and their target genes in mice were retrieved from Regulatory Network Repository (http://www.regnetworkweb.org/) and Transcriptional Regulatory Element Database (TRED; http://www.cb.utdallas.edu/cgi-bin/TRED) and their association with leishmaniasis was done through extensive literature survey. The data, thus derived, was also matched to RNA-seq data that dealt with comparison of *L. major* infection in peritoneal macrophages at different time points. In total, 71 genes were incorporated in the network, which has a bipartite architecture. This network was then put through node selection in Cytoscape (Version 3.4.0) by considering each nodes closeness centrality and edge betweeness.

### Multi Objective Optimization and Evolvability

Evolvability of a biological system is defined as the amenability for evolutionary changes. Biological evolvability depends upon how malleable the genotype is to phenotypic mapping that creates morphological diversification. Since evolvability is a multi-objective optimization problem, the system under consideration was put through a multi-objective optimization. Optimization also helps in the bioengineering of novel signaling/metabolic pathways using synthetic biology approach of the natural system (optimization). The solution to the multi objective functions are a set of optimal solutions known as the Pareto optimal solutions. For our built-in TFTG network, optimization was achieved by defining two opposing objective functions; one that considers the system associated with anti-inflammatory condition and the other system associated with pro-inflammatory condition (**Figure 2**). The multi-objective optimization ^33^ was performed in MATLABs’ Optimization toolbox (7.11.1.866) (The MathWorks Inc.).

**Figure 2:**
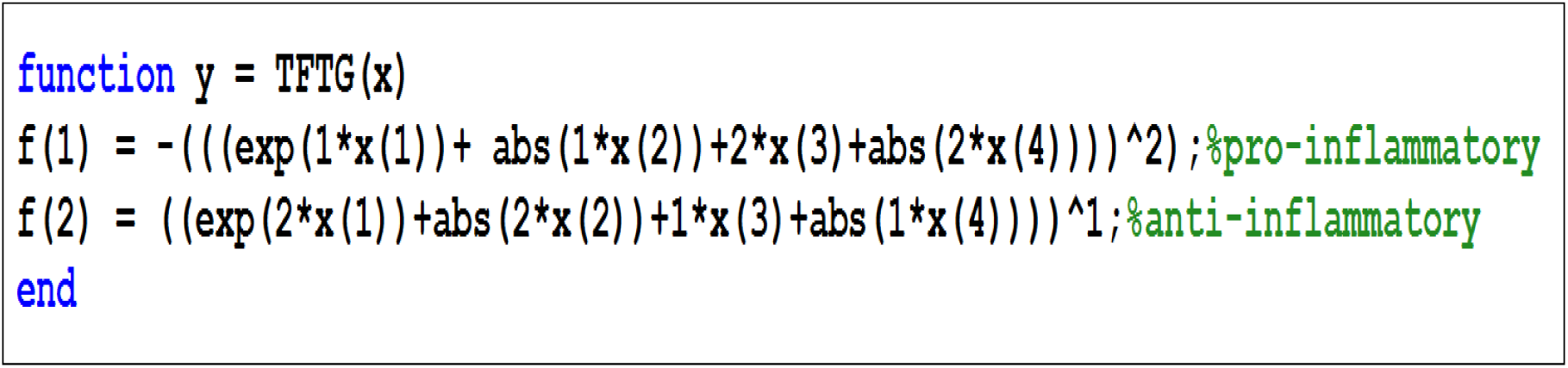
The objective functions defined for the TFTG evolvability: f(1) function depicting the pro-inflammatory and f(2) function depicting the anti-inflammatory condition.

The two objective were implemented using the solver “gamultiobj” and the parameters set for the algorithm is given in **Table 1**

**Table 1:**
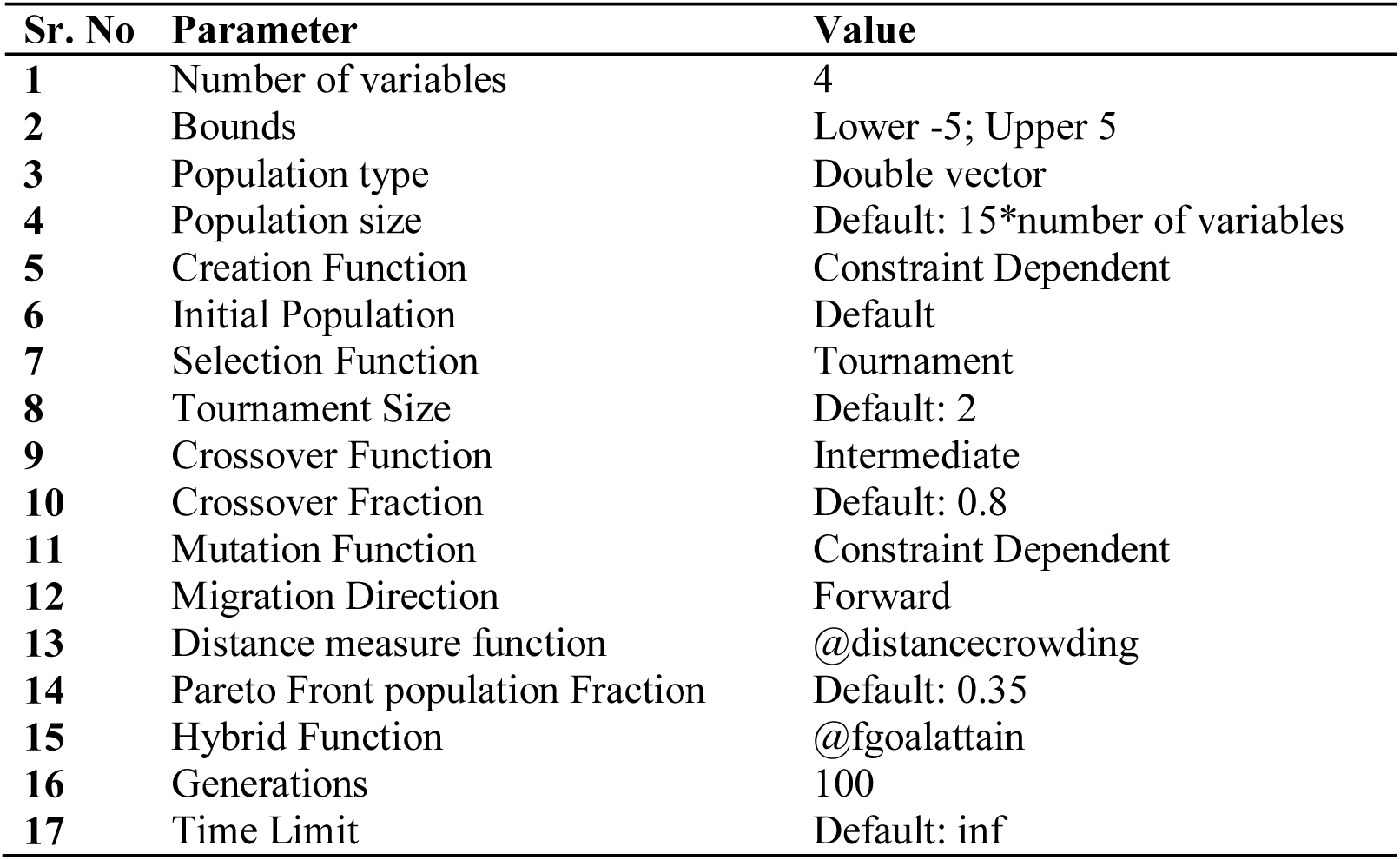
Parameters set for the implementing “gamultiobj”.

| Sr. No | Parameter | Value |
| --- | --- | --- |
| 1 | Number of variables | 4 |
| 2 | Bounds | Lower -5; Upper 5 |
| 3 | Population type | Double vector |
| 4 | Population size | Default: 15*number of variables |
| 5 | Creation Function | Constraint Dependent |
| 6 | Initial Population | Default |
| 7 | Selection Function | Tournament |
| 8 | Tournament Size | Default: 2 |
| 9 | Crossover Function | Intermediate |
| 10 | Crossover Fraction | Default: 0.8 |
| 11 | Mutation Function | Constraint Dependent |
| 12 | Migration Direction | Forward |
| 13 | Distance measure function | @distancecrowding |
| 14 | Pareto Front population Fraction | Default: 0.35 |
| 15 | Hybrid Function | @fgoalattain |
| 16 | Generations | 100 |
| 17 | Time Limit | Default: inf |

**Table 2:**
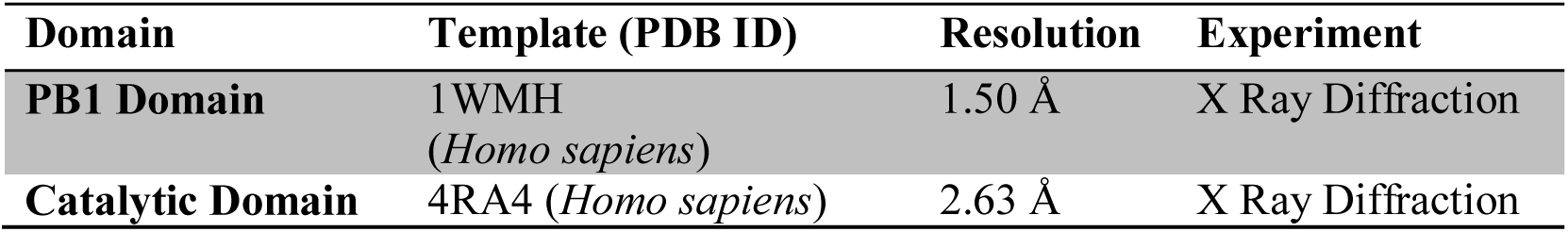
The PDB IDs of templates used for homology modeling of the chimeric PKC_ζα.

| Domain | Template (PDB ID) | Resolution | Experiment |
| --- | --- | --- | --- |
| **PB1 Domain** | 1WMH<br>(*Homo sapiens*) | 1.50 Å | X Ray Diffraction |
| **Catalytic Domain** | 4RA4 (*Homo sapiens*) | 2.63 Å | X Ray Diffraction |

**Table 3:** CT_PKC parts with the corresponding Parts registry catalogue number.

| Sr. No | Part | Number of bp | Parts registry catalogue number |
| --- | --- | --- | --- |
| A | lacO2 operator | 20 | BBa_K731500 |
| B | LacI Repressor Protein | 1086 | BBa_K731500 |
| C | P2A Sequence | 66 | BBa_K1537016 |
| D | GSG linker sequence | 9 | Ref() |
| E | Chimeric _PKC | 1392 | Designed in house |
| F | T2A Sequence | 63 | BBa_K1537017 |
| G | GFP Reporter protein | 1257 | BBa_K194002 |
| H | Terminator Sequence | 41 | BBa_B0012 |

### Chimeric PKC design

The chimeric PKC was designed using the concept of evolutionary domain shuffling. It is one the major mechanisms leading to the formation of new proteins during the course of evolution. As evolutionary domain shuffling is associated with emergence of novel biological function, synthetic domain shuffling can be applied to have proteins engineered with a novel cellular function. Chimeric PKC (462 amino acids) was designed by taking the PB1 domain and catalytic domain from two isoforms of PKC (**Figure 3**). The N terminal of chimeric PKC protein was tagged with 6XHIS.

**Figure 3:**
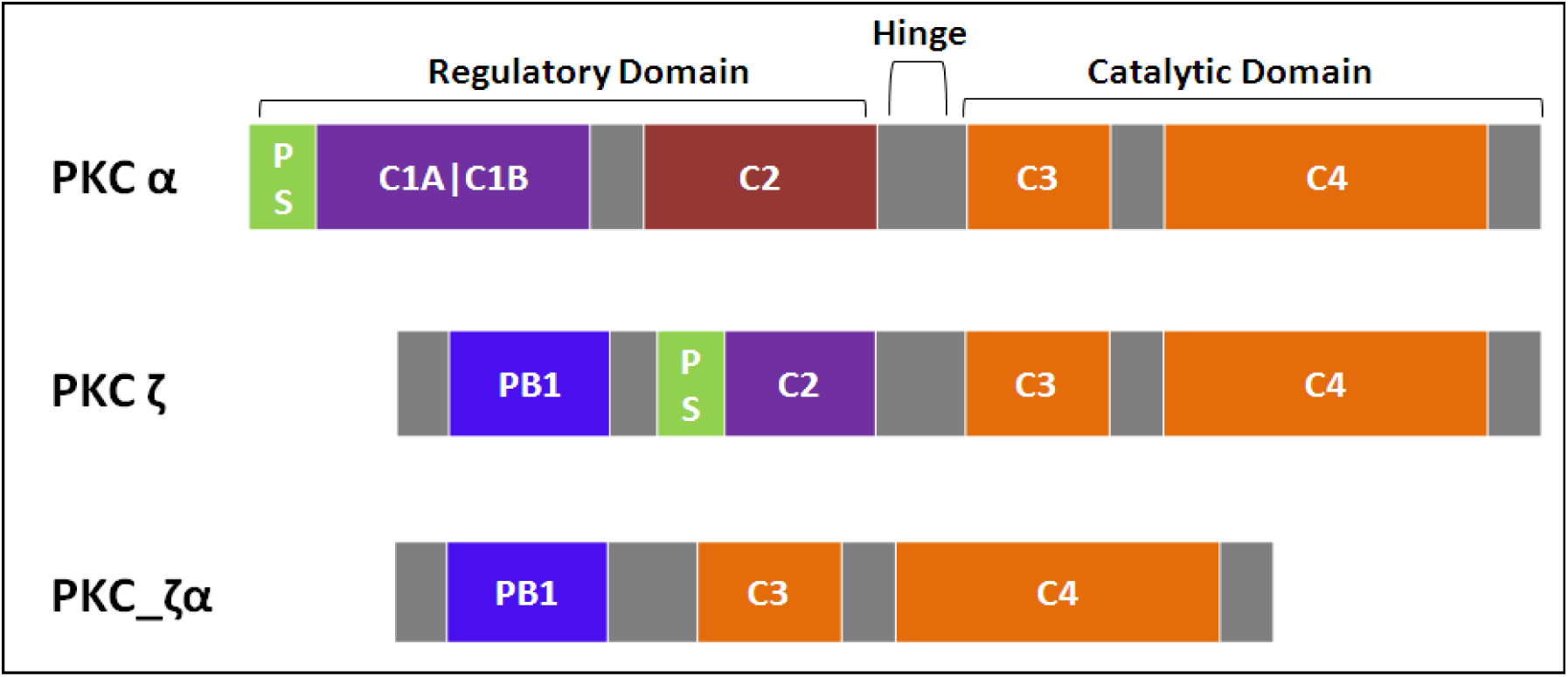
Chimeric PKC design derived from the conventional and atypical isoforms of PKC.

The amino acid sequence details are as follows

#### PBI Domain from PKC_ζ(Atypical PKC)

Domain one: PB1 (16 – 98 aa)

>gi|84872200:16-98 protein kinase Cζ type isoform [*Mus musculus*]

VRLKAHYGGDILITS VDAMTTFKDLCEEVRDMCGLHQQHPLTLKWVDSEGDPCTVS SQMELEEAFRLVCQGRDEVLIIHVFPS

#### Catalytic Domain from PKCα (Conventional PKC)

Domain two: Catalytic (290 – 668 aa)

>gi|164663791:290-668 protein kinase C α type isoform [*Mus musculus*] IPEGDEEGNMELRQKFEKAKLGPAGNKVISPSEDRKQPSNNLDRVKLTDFNFLMVLG KGSFGKVMLADRKGTEELYAIKILKKDVVIQDDDVECTMVEKRVLALLDKPPFLTQ LHSCFQTVDRLYFVMEYVNGGDLMYHIQQVGKFKEPQAVFYAAEISIGLFFLHKRGI IYRDLKLDNVMLDSEGHIKIADFGMCKEHMMDGVTTRTFCGTPDYIAPEIIAYQPYG KSVDWWAYGVLLYEMLAGQPPFDGEDEDELFQSIMEHNVSYPKSLSKEAVSICKGL MTKHPAKRLGCGPEGERDVREHAFFRRIDWEKLENREIQPPFKPKVCGKGAENFDKF FTRGQPVLTPPDQLVIANIDQSDFEGFSYVNPQFVHPIL

### Homology modeling and MD simulation of the chimeric PKC

Since the chimeric PKC is a synthetically engineered protein, its 3D structure was modeled from well understood crystal structure of the ζ and α PKC isoforms, using homology or comparative modeling in Modeller (v 9.11) ^34^. Comparative modeling consists of four main steps (i) template(s) identification (ii) alignment of the target and the template sequence (iii) model building i.e. assigning coordinates to the target sequence with respect to the chosen target, and finally (iv) predicting model errors ^34^. The template for the chimeric PKC was chosen by performing protein BLAST (pBLAST). Protein sequences with highest sequence similarity and identity where selected as templates for chimeric PKC_ζα. The .PDB format for the crystal structures of the templates where retrieved from RCSB PDB (Research Collaboratory for Structural Bioinformatics-Protein Data Bank) (www.rcsb.org).

The target and template sequences were aligned in the .PIR (Protein Information Resource) format in MultAlin ^35^. The .PIR alignment file of the target and template and the .PDB file of the templates are used as the input file in Modeller. The model that was built in Modeller had many outliers which were refined using the loop refinement or optimization protocol. The final model was then analyzed with ProSA-Web ^36^ and RAMPAGE ^37^ tool for model quality and check for any errors pertaining to the built 3D structure of the chimeric PKC_ζα the figures for the modeled chimeric PKC_ζα were generated in PyMOL ^38^ Molecular Graphics System, Version 1.7.4.5 Schrodinger, LLC.

Stability analysis under physiological condition of the chimeric PKC_ζα logy model was performed by molecular dynamic OPLS 2005 simulation in DESMOND 3.2 (D.E. Shaw Research) from Maestro 8.2 ^39^. Transferable intermolecular potential with 3-points (TIP3P) water model was used in the orthorhombic box with a default 10nm cut-off periodic boundary condition (PBC). Default long range and short range parameters were used along with a 2000 round of steepest descent minimization and Berendsen Normal Pressure temperature/ Normal Volume Temperature (NPT/NVT) equilibration was performed. After equilibration a production MD run was performed with a time step of 2fs and for a time scale of 15ns. All essential parameters such as potential energy, pressure, temperature were checked throughout the length of the MD simulation. The root mean square deviation (RMSD), root mean square fluctuation (RMSF) and other parameters of the trajectories were also calculated using default Desmond event analysis protocols and minimum energy structures were exported from the trajectories for further analysis.

### Synthesis, expression and identification of the chimeric PKC

The amino acid sequence for the chimeric PKC_ζα protein was given for synthesis to GeneArt, Invitrogen. The sequence was reverse translated (1422 bp) and codon optimized for *E. coli* expression system. The synthesized nucleic acid was inserted into the pET151 Topo vector. The vector with the insert was digested using restriction enzymes EcoRI and HindIII for the confirmation of the insert. The construct was transformed into competent BL21 (DE3) by heat shock treatment at 42°C for 30 sec. These cells were plated on ampicillin containing Luria Berttini (LB) agar plates and incubated at 37°C overnight. The transformed cells (ampicillin resistant) were selected and used for expression of the chimeric PKC. The expression of the protein was induced by 1mM IPTG at 37°C for 1 hour and continued till 3 hours. The expression of the chimeric PKC_ζα was checked on SDS-PAGE and analyzed by LC-MS/MS. In gel sample for MS analysis was prepared by cutting the expressed protein from the gel with a razor and thoroughly rinsing the gel piece with the coomassie stained gel 25 mM ammonium bicarbonate (ABC). The gel is dehydrated and rehydrated with alternate washes of 2:1 mixture of Acetonitrile (ACN): 50 mM ABC and 25 mM ABC. After which the gel is alkylated with 10-20 mM Dithiothreitol (DTT) followed by 50-100 mM Iodoacetamide (IAA). Again the gel is cycled through the dehydration and hydration and digested with trypsin (Sigma/Promega MS grade trypsin). The gel is extracted using solution (50:50 ACN: H2O, 0.1% Trifluoroacetic Acid (TFA)), and the supernatant is completely dried in a Speed Vac and later resuspended in 10 μL of 0.1 % TFA in water and used for MS analysis at the Proteomics Facility, NCCS, Pune.

### Western Blotting

Western blot was done to show the phosphorylation of IkkB by the chimeric PKC_ζα on transfection followed by induction with 1mM IPTG. Peritoneal macrophages were isolated from Balb/c mice, post four days of injection of 3% thioglycolate media into the peritoneal cavity of the mice. These cells (2X10^6^ cells/ml) were plated and allowed to adhere to the culture vessel for 2-3 hours at 37°C in the CO2 incubator at 5% humidity. The non-adherent cell were washed with warm PBS and the cells where transfected with chimeric PKC_ζα construct using the Lipofectamine 3000^®^ (Invitrogen) following the manufacturer’s protocol. Briefly, Lipofectamine 3000 reagent was diluted in Opti-MEM (Gibco, Life Technologies^®^) media and the PKC construct (DNA) (5μg) was diluted in Lipofectamine 3000 reagent and Opti-MEM media. Diluted Lipofectamine reagent and DNA were mixed in the ratio of 1:1 and kept at room temperature for 5-10 minutes to allow the formation of DNA-lipid complex. The DNA-lipid complex was added to the peritoneal macrophages and incubated for 6hrs at 37°C in the CO2 incubator. After the initial incubation, the plates are replenished with fresh Roswell Park Memorial Institute (RPMI) 1640 supplemented with 10% fetal bovine serum (FBS) (Sigma^®^). The transfected cells were induced with 1mM IPTG for 24 hours, followed by cell lysis, lysate collection, SDS-PAGE and western blotting. Densitometric analyses for the bands were carried out using ImageJ. Band intensities were quantified and the values were normalized to endogenous control β-actin and expressed as relative densities, indicated as bar graphs adjacent to corresponding figures.

### Confocal Microscopy

To show that the chimeric PKC _ζα indeed causes the IkkB phosphorylation, confocal microscopy was performed. Briefly, peritoneal macrophages were plated at 2x10^5^ cell/ml in 8 welled chamber slides and were transfected with the chimeric PKC_ζα construct with Lipofectamine 3000^®^ (Invitrogen) reagent. These cells were induced with 1mM IPTG for 24hrs and prepared for confocal microscopy. 4% paraformaldehyde (PFA) was used for fixation, followed by permeabilization in 0.1% Triton X. Blocking was done in 5% normal goat serum/2% normal horse serum, after which cells were probed with primary antibody against NFkB at room temperature (RT) for 60 mins. After washing with PBS secondary antibody Alexa Flour 594 (a kind gift from Dr. Jomon Joseph, NCCS, Pune) was added to the cells and incubated for 30 mins at RT followed by washing with PBS. The slides were dried and mounted with VectaShield mounting medium (Vector Laboratory ^®^) and observed under Zeiss LS 600 confocal microscope, NCCS Confocal facility.

### Negative autoregulatory circuit design and its quasipotential landscape

The synthetic circuit with a LacI negative autoregulatory loop was designed in Tinker cell (www.tinkercell.com) and the chimeric PKC_ζα was coupled to it in such a way that its expression was under the transcriptional control of LacI repressor protein. Two ODEs depicting the negative autoregulatory structure of the circuit was written in Berkeley Madonna (v8.3.18) (www.berkeleymadonna.com) and simulated using the Runge Kutta 4 (RK4) method. Time series data for 100s was generated and analysed by Gene Regulatory Network Inference sing Time Series (GRENITS) (v1.24.0) ^40^ and BoolNet (v2.1.3) ^41^ packages in Bioconductor’s R console. Circuit’s convergence, possible attractor states the circuit can be found in, the probability of Boolean network transitions and network wiring were created. Nullclines for the system of ODEs was also generated in Berkeley Madonna. The quasipotential landscape depicting the switching mechanism was inferred using the equation Vq= -((LacR)+(PKC))*DT derived from the Waddington’s evolutionary landscape^42^.

### *In vitro* verification of the working of the synthetic circuit

The designed synthetic construct with a negative autoregulatory loop referred to as CT_PKC was synthesized by GeneArt, Invitrogen^®^. The order of the parts used in the design CT_PKC is mentioned below:

The parts were placed downstream of the CMV promoter in pcDNA 3.1(+). The synthesized construct was then analyzed by restriction digestion with BamHI.

The construct was transfected into peritoneal macrophages by Lipofectamine 3000^®^ reagent (Invitrogen) according to manufacturer’s protocol. After 24hrs, transfection efficiency was calculated and was found to be around 60%. The working of the construct was ascertained, by induction using 1 mM IPTG for 24 hours and looking for GFP expression under EVOS, Life Technologies, fluorescence microscope.

### *In vitro* effect of cytokine gene expression and NO as a result of action of chimeric PKC_ζα and miltefosine

To check the efficacy of the chimeric PKC_ζα in combination with miltefosine and *L. major* infection, on cytokine gene expression, quantitative PCR analysis was performed. Peritoneal macrophages were used for seven treatment groups; namely Infection (I), Chimeric PKC (CT), Chimeric PKC + Infection (CTI), Chimeric PKC + Miltefosine (CTM) and Chimeric PKC + Infection + Miltefosine (CTIM), which were compared to the untreated control.

For the infection groups, peritoneal macrophages were infected with stationary phase 2X10^7^ and 1X10^7^ promastigotes at a cell to parasite ratio of 1:10 to 1:5 respectively. After 16 hours, of incubation with the parasite the cells were extensively washed with PBS to remove adherent extracellular parasites. The infectivity index calculation was performed by counting the number of intracellular parasites after DAPI staining using EVOS florescence microscope (60X magnification).

Miltefosine treatment was used at a concentration of 5uM which is non-cytotoxic to peritoneal macrophages. The transfected cells were induced with 1mM IPTG. RNA samples were collected at time intervals of 0, 24, 48 and 72 hours from all the treatment and control groups. cDNA was prepared by reverse transcription using the High Capacity cDNA Reverse Transcriptase Kit^®^, Applied Biosystems using the cycling conditions as given below:

| **Settings** | **Step1** | **Step2** | **Step3** | **Step4** |
| --- | --- | --- | --- | --- |
| **Temperature** | 25 °C | 37 °C | 85 °C | 4 °C |
| **Time** | 10 minutes | 120 minutes | 5 minutes | ∞ |

This cDNA was used as template for qPCR assay using the Taqman primer probe (FAM - 6-carboxyfluorescein) mix listed below:

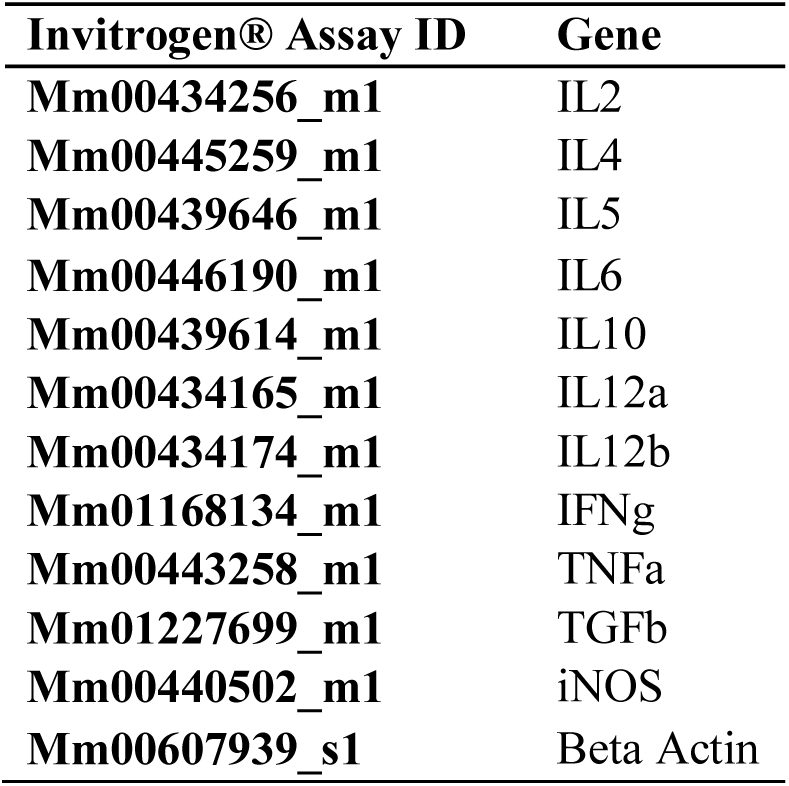

| **Invitrogen® Assay ID** | **Gene** |
| --- | --- |
| **Mm00434256_m1** | IL2 |
| **Mm00445259_m1** | IL4 |
| **Mm00439646_m1** | IL5 |
| **Mm00446190_m1** | IL6 |
| **Mm00439614_m1** | IL10 |
| **Mm00434165_m1** | IL12a |
| **Mm00434174_m1** | IL12b |
| **Mm01168134_m1** | IFNγ |
| **Mm00443258_m1** | TNFα |
| **Mm01227699_m1** | TGFβ |
| **Mm00440502_m1** | iNOS |
| **Mm00607939_s1** | Beta Actin |

The cycling condition for qPCR on the StepOnePlus™ Real-Time PCR System is given below

| **Settings** | **Step1** | **Step2** | **Step3** | **Step4** |
| --- | --- | --- | --- | --- |
| **Temperature** | 50°C | 95°C | 95°C | 60°C |
| **Time** | 2 minutes | 20 sec | 0.01 sec | 0.30 sec |
| **Condition** | Hold |  | Cycle (40 Cycles) |  |

Nitrite estimation was done using the Greiss’s method (Greiss’s Kit from Invitrogen^®^)

### Statistical Analysis

All the experiments have been done in triplicates and one way ANOVA with Tukey’s correction was employed for statistical significance.

### Animal Ethics

The Balb/c mice used for peritoneal macrophage collection were placed in the NCCS, Experimental Animal Facility (7/GO/c/99/CPCSEA), and all norms for the ethical use of animals were stringently followed (Project No.B-198).

## 3. Results

### Reconstruction of TFTG network and Node selection

The whole TFTG network was analysed using the network analyser in Cytoscape (Version 3.4.0) and the results are tabulated in **Table 4**. The TFTG network is made up of 134 transcription factor and target gene pairs, from which important TFs and their TG genes were selected based on network matrices of closeness centrality (CC) and edge betweeness. CC measures the reciprocal of farness of a node in a network i.e. higher the value of CC more important is the node in the network. The importance of the node is due the fact that higher the CC faster will be information spread to the reachable nodes. In the reconstructed network, NFkB shows the highest CC value (**Figure 4(a) and (b)**), indicating that this TF is an important node in transmitting signaling information for appropriate gene expression. From the node pair list and the network matrices applied to the TFTG network, it can be seen that cytokine genes like IL2, IL4, IL5, IL6, IL10, IL12b, Nos2 and Tnfa can be affected the most, if NFkB is modulated.

**Table 4:**
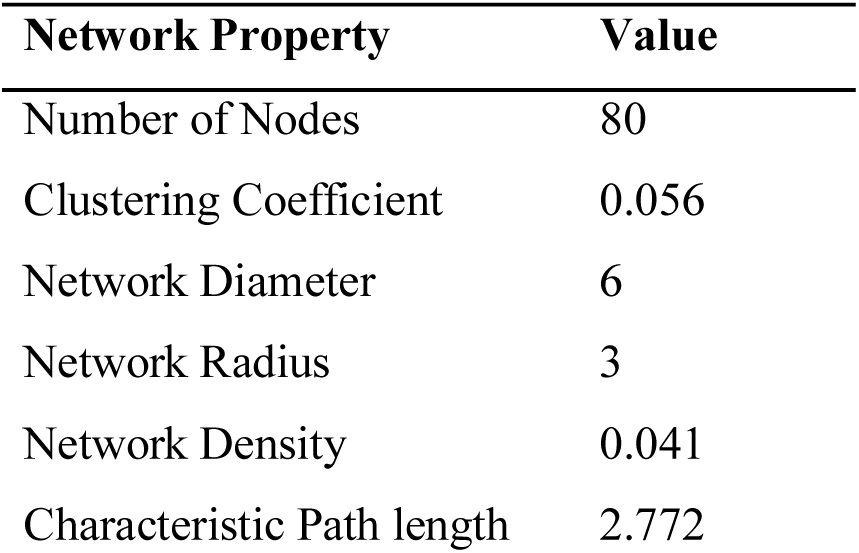
Whole TFTG network property.

**Figure 4:**
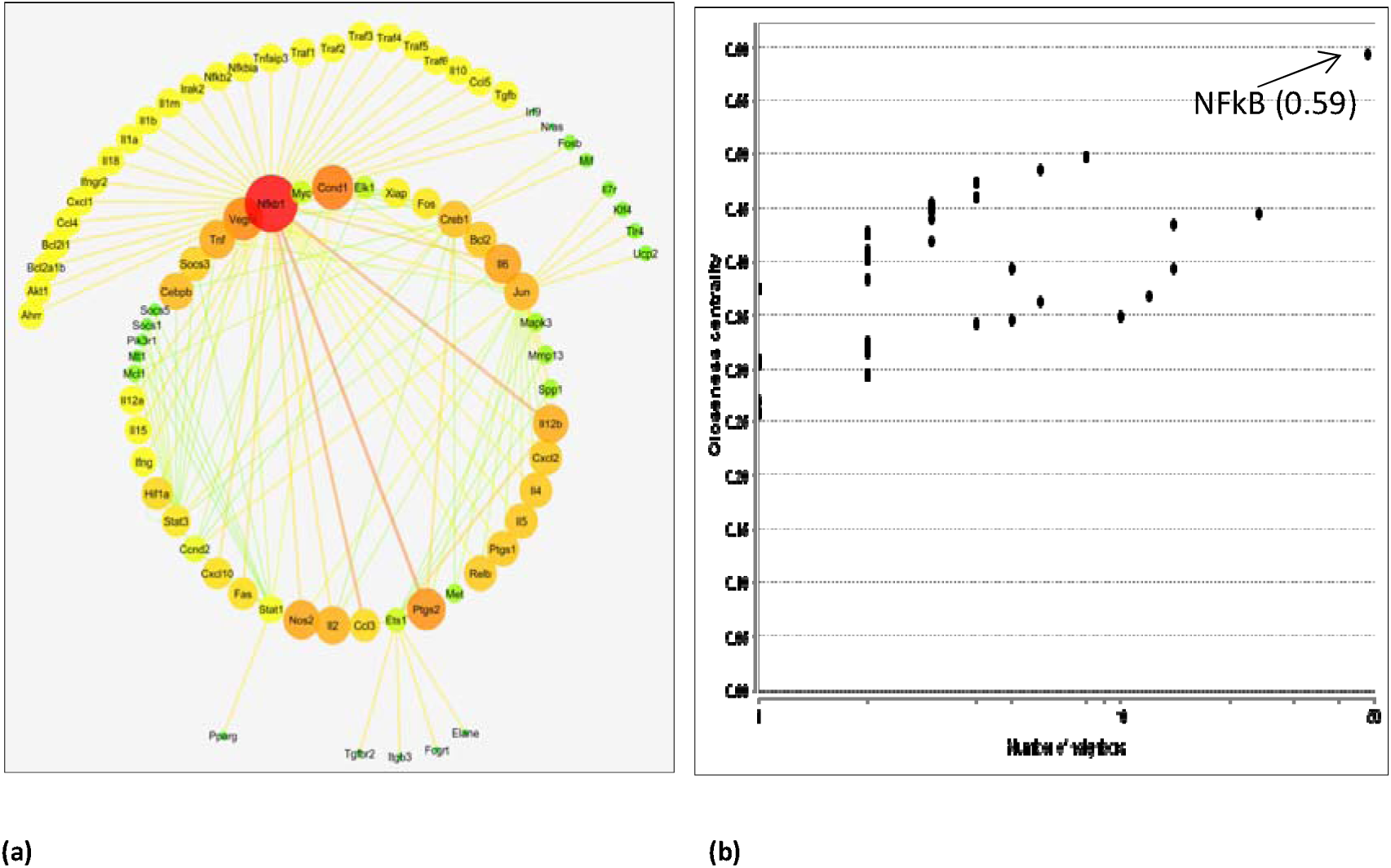
(a) Node selection based on closeness centrality and edge betweeness (b) Closeness centrality values of all the nodes in the network and NFkB1 showing the highest CC value of 0.59.

### Optimization and Evolvability of the reconstructed TFTG network

The objective function defined and solved by the “gamultiobj” solver shows that the system is evolvable to a new phenotype. As cited earlier the NSGA-II algorithm, during the selection procedure, the best offspring population is combined with the current generation population. This step ensures that elitism is maintained during the run of the genetic algorithm (**Figure 5 (a)**). The population is then sorted based on non-domination, and this procedure is continued till the population size exceeds the current population size to generate the subsequent generations. The score histogram for the defined objective functions shows that f(2) is higher compared to the f(1), but the number of individuals i.e. solutions to the given objective function lie more in favour of the f(1). The f(1) function defines the pro-inflammatory phenotype of the system (**Figure 5 (b)**). The Pareto fronts (**Figure 5 (c)**) for the opposing objective functions have 19 non-dominated solutions in a single run and the average spread measure for these non-dominated solutions is 0.129942 (**Figure 5 (d)**).

**Figure 5:**
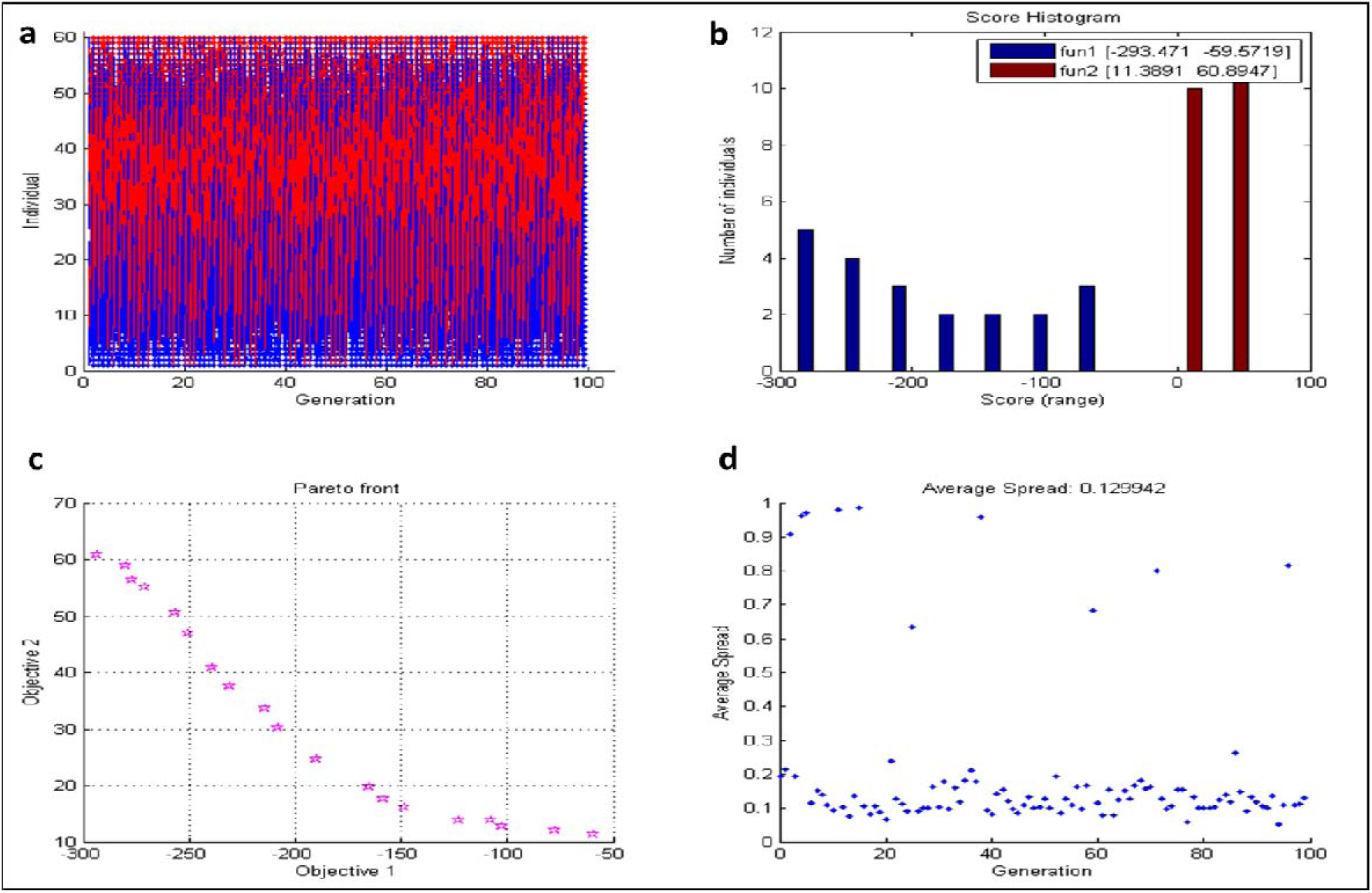
Evolvability Analysis of the TFTG network.

The decision variables for the multiobjective function were the cytokine, whose concentration would be measured in the *in vitro* experiments. They are denoted as x1, x2, x3 and x4, representing the cytokines IL10, IL4, TNFa and IFNg respectively. These decision variables are embedded in the f1 and f2 objective functions. The fitness value obtained for the objective function decides the trend that is followed by the decision variable as depicted in **Figure 6.** It shows that the x1 and x2 decision variables depicting the anti-inflammatory cytokines show a rugged oscillatory behaviour, converging to the f2 fitness function. Similarly, x3 and x4 depicting pro-inflammatory converges towards the f1 fitness function. The overall fitness landscape for the two objectives shows a convergence to f1 fitness (**Figure 7**), which is the ultimate goal of the multi-objective optimization in this study.

**Figure 6:**
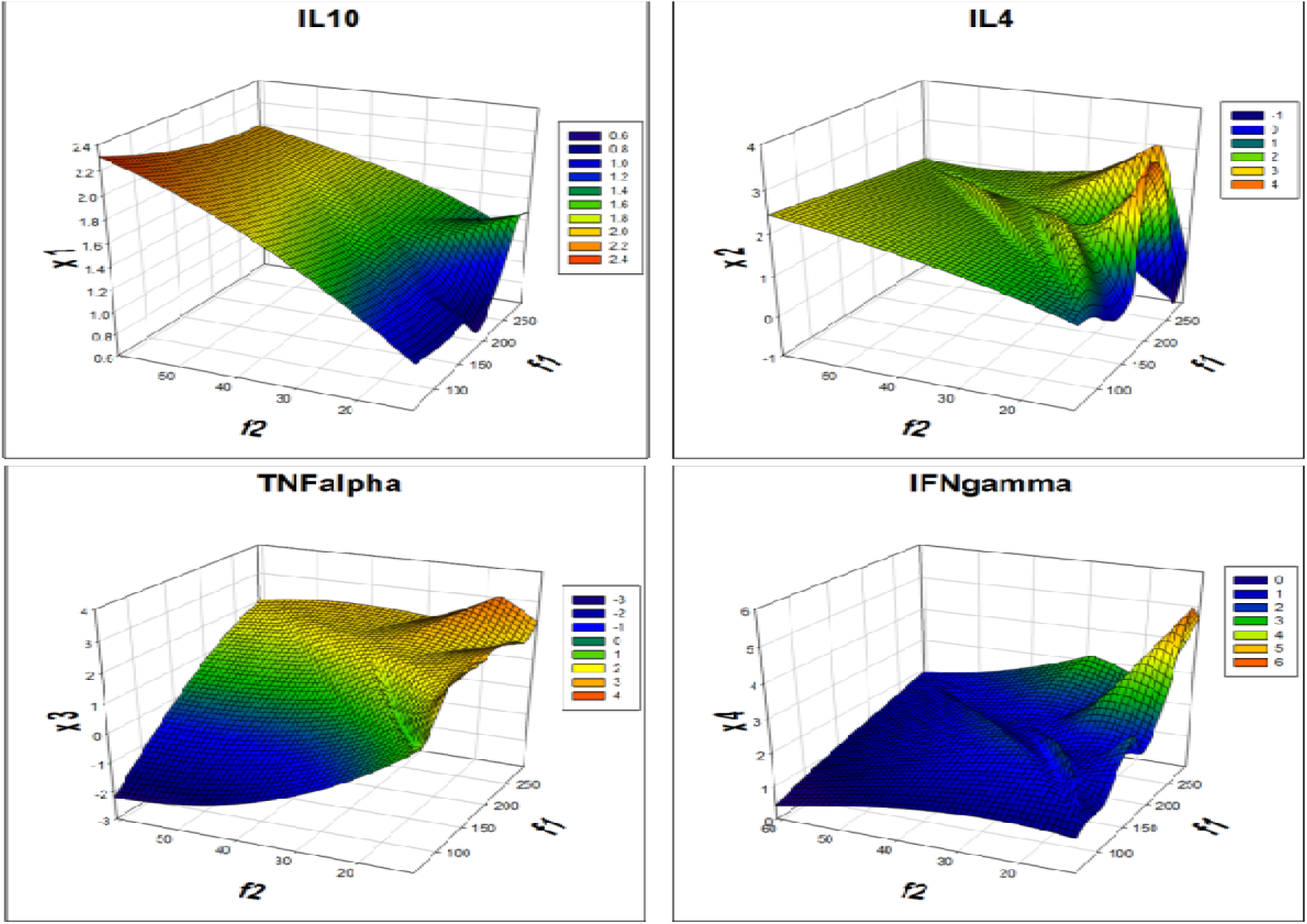
Fitness value V/s decision variable for the anti-inflammatory cytokines IL10 and IL4 and pro-inflammatory cytokines TNFalpha and IFNgamma.

**Figure 7:**
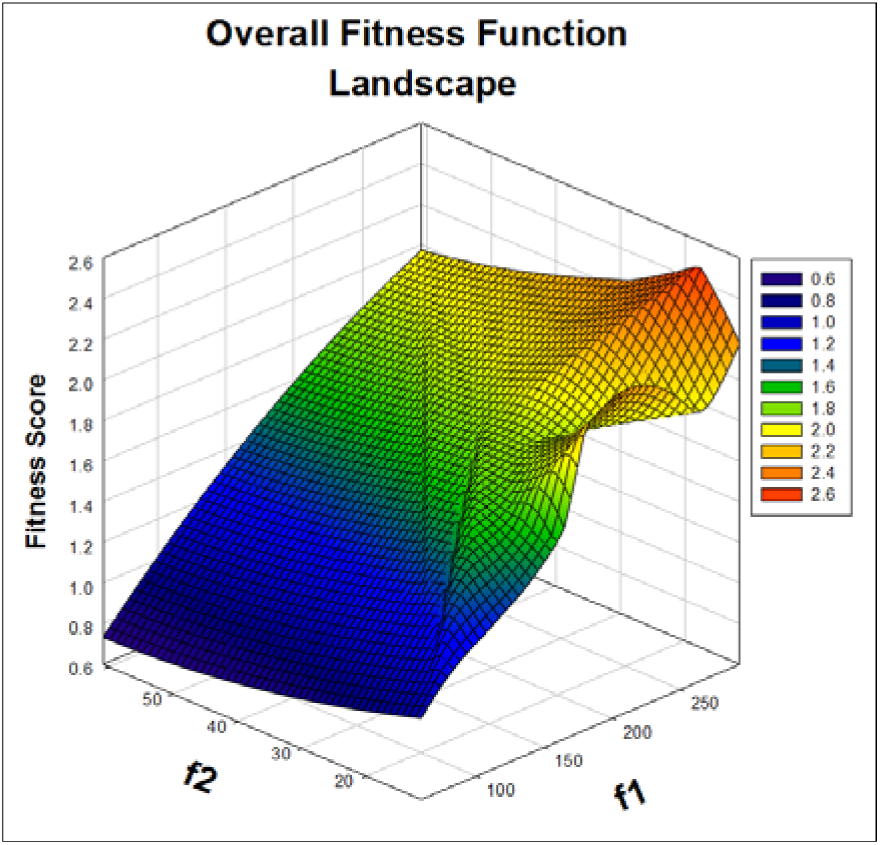
The overall fitness function landscape for the objective function f1 and f2.

### Chimeric PKC, its Homology Modeling and MD simulation

The sequence of amino acid of the chimeric PKC_ζα was analyzed by the ProtParam ^43^ tool for the physiochemical properties, which are listed below.

Molecular weight: 52626.4 Da

Theoretical pI: 5.26

Amino acid composition:

| Aa | % | Aa | % | Aa | % | Aa | % |
| --- | --- | --- | --- | --- | --- | --- | --- |
| Ala | 4.5% | Leu | 8.9% | Gln | 4.3% | Ser | 4.3% |
| Arg | 4.3% | Lys | 7.4% | Glu | 8.0% | Thr | 3.7% |
| Asn | 2.6% | Met | 3.5% | Gly | 7.4% | Trp | 0.9% |
| Asp | 7.6% | Phe | 5.8% | His | 2.8% | Tyr | 3.0% |
| Cys | 2.4% | Pro | 5.6% | Ile | 5.6% | Val | 7.4% |

Instability index: The instability index (II) is computed to be 36.47, classifying the protein as stable.

Aliphatic index: 82.45

Grand average of hydropathicity (GRAVY): −0.316

Homology modeling of the chimeric PKC_ζα was performed using 1 WMH and 4RA4 as templates. The homology model of chimeric PKC_ζα, shows about 96.5% of amino acid residues in the favored region and 2.6% in the allowed region. Eleven amino acids were in the outlier region and so loop refinement of the structure was done after which the number of outlier amino acids reduced to 4 i.e. 0.9% (**Figure 11 (a)**). This model was used as the starting model for MD and the protein shows stable dynamics over the 15ns simulation. All the physical parameters viz temperature, volume and pressure were maintained during the 15ns MD run (**Figure 8**). RMSD plot of 15ns MD simulation for showing the mean fluctuation was at 8 Angstrom and stabilized after 10ns while RMSF plot of the 462 amino acids of the Chimeric PKC_ζα shows that for apart from the hinge/loop of the chimeric PKC all the other amino acids show a mean fluctuation within the 1 Angstrom range (**Figure 9 (a) and (b)**).

**Figure 8:**
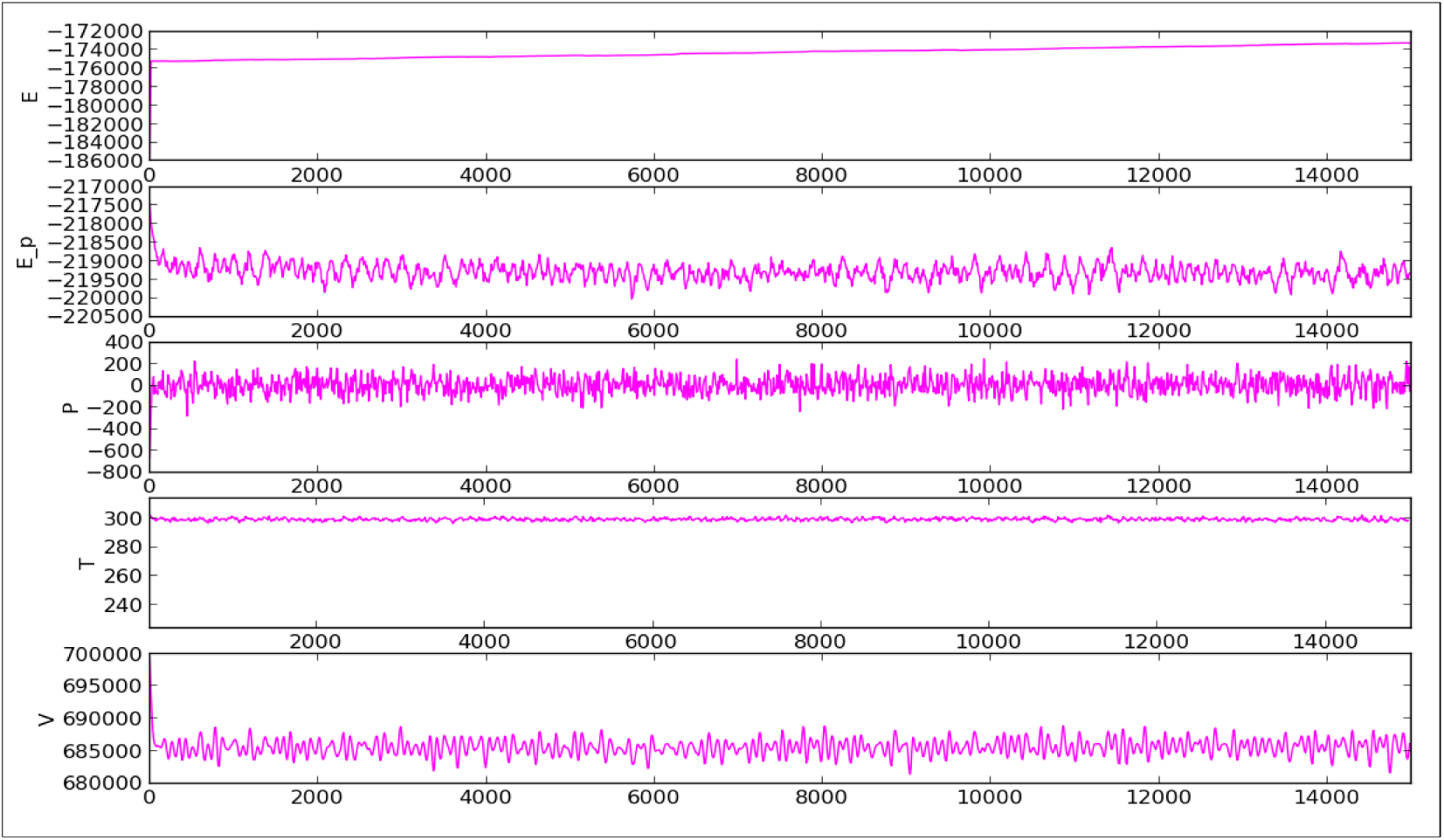
Physical parameters during the MD simulation were constant throughout the simulation time of 15ns.

**Figure 9:**
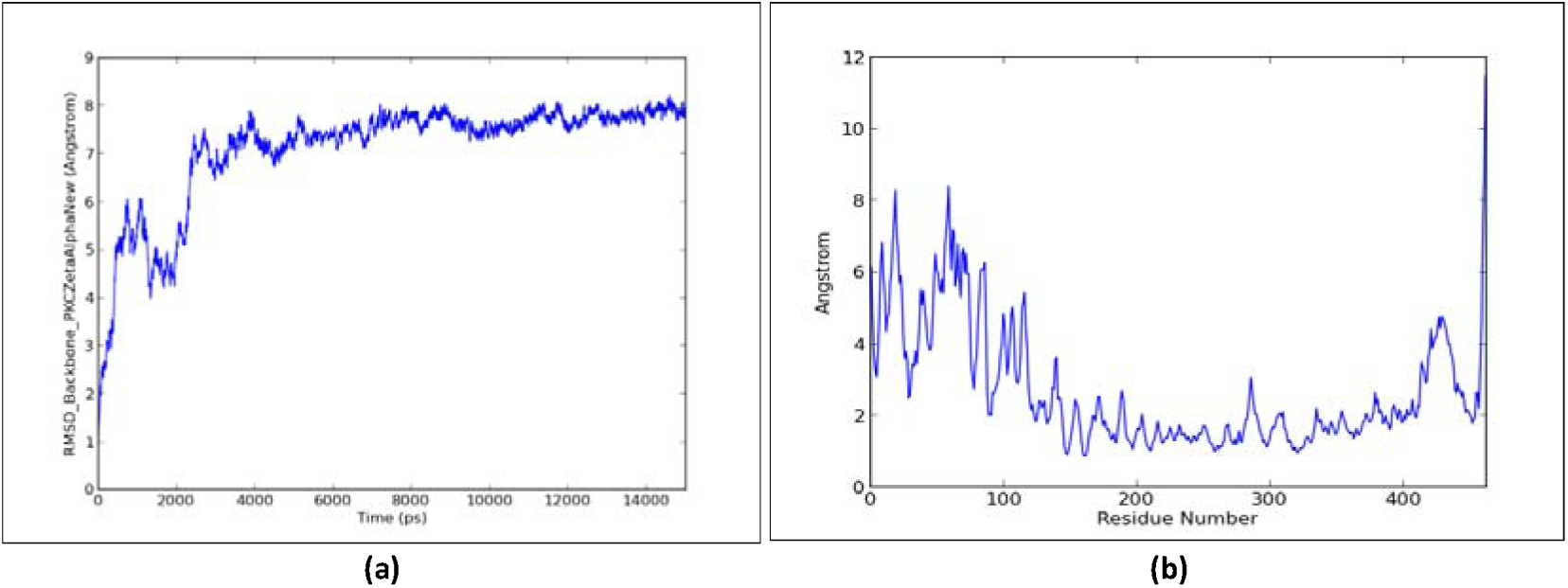
(a) RMSD plot of 15ns MD simulation for showing the mean fluctuation was at 8 Angstrom and stabilized after 10ns; (b) RMSF plot of the 462 amino acids of the Chimeric PKC_ζα.

To check for any modelling errors, the 3D model obtained from modeller and the 3D model after MD simulation was checked on ProSa-Web which returns the z-score and energy plots for the chimeric PKC (**Figure 10**). The z-score indicates overall model quality and any deviation in the energy structure from random conformations. Z-scores of the chimeric PKC is well within the z-scores of all the experimentally (X-ray, NMR) determined protein structures in PDB. Similarly, the energy plot shows the local model quality by plotting energies as a function of amino acid sequence at a particular position. If most of the amino acids in the structure show a negative energy value then it can be said that the structure is error free, which is the case with the chimeric PKC. The final 3D structure after MD simulation of the chimeric PKC is depicted in **Figure 11 (b).**

**Figure 10:**
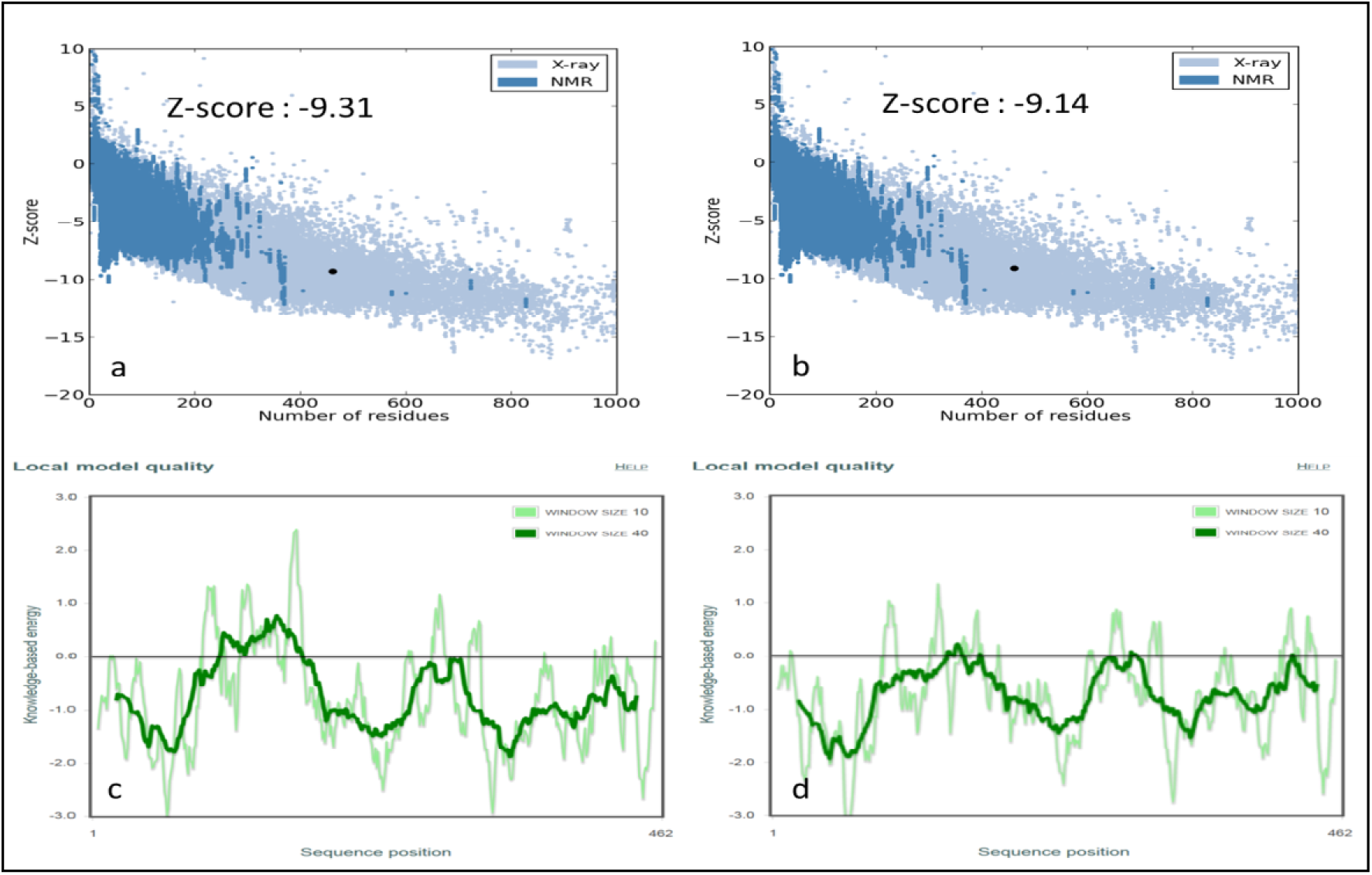
ProSA-Web Analysis of the chimeric PKC_ζα before and after MD simulation: a and c before MD and b and d after MD (a and b depicts the Z-score; while c and d depict the energy plot form each amino acid)

**Figure 11:**
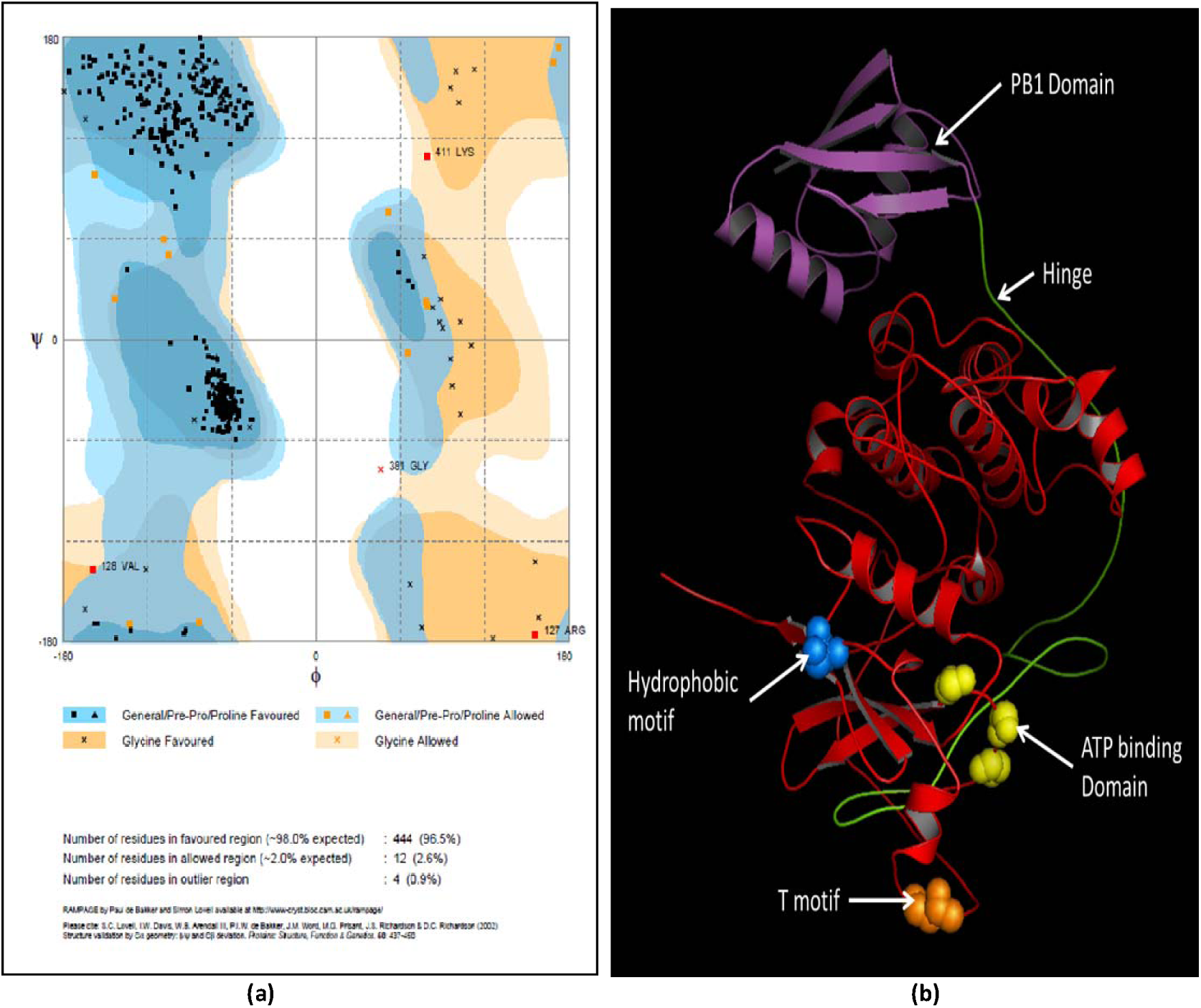
(a) Rampage Analysis of chimeric PKC_ζα (Number of residues in favoured region (‘98.0% expected): 444 (96.5%); Number of residues in allowed region (‘2.0% expected) : 12 (2.6%); Number of residues in outlier region: 4 (0.9%)) (b) Final model of the chimeric PKC_ζα after MD showing the conserved regions; ATP binding domain (yellow): MVLG^140^KG^142^SFG^14S^; T motif (orange): PVLT^431^PPDQL; Hydrophobic motif (blue): DFEGFS^450^YVNPQ; Purple is the PB1 domain and Red is the catalytic domain.

### Synthesis, expression and identification of the chimeric PKC

The construct was analyzed by restriction digestion and confirmed the presence of the insert (**Figure 12**). The chimeric PKC was successfully transformed in to BL21DE3 and induced for the expression of chimeric PKC (**Figure 13 (a)**) which was confirmed by MS analysis. The His-tagged chimeric protein was also confirmed by western blotting using the anti-His antibody (**Figure 13 (b)**).

**Figure 12:**
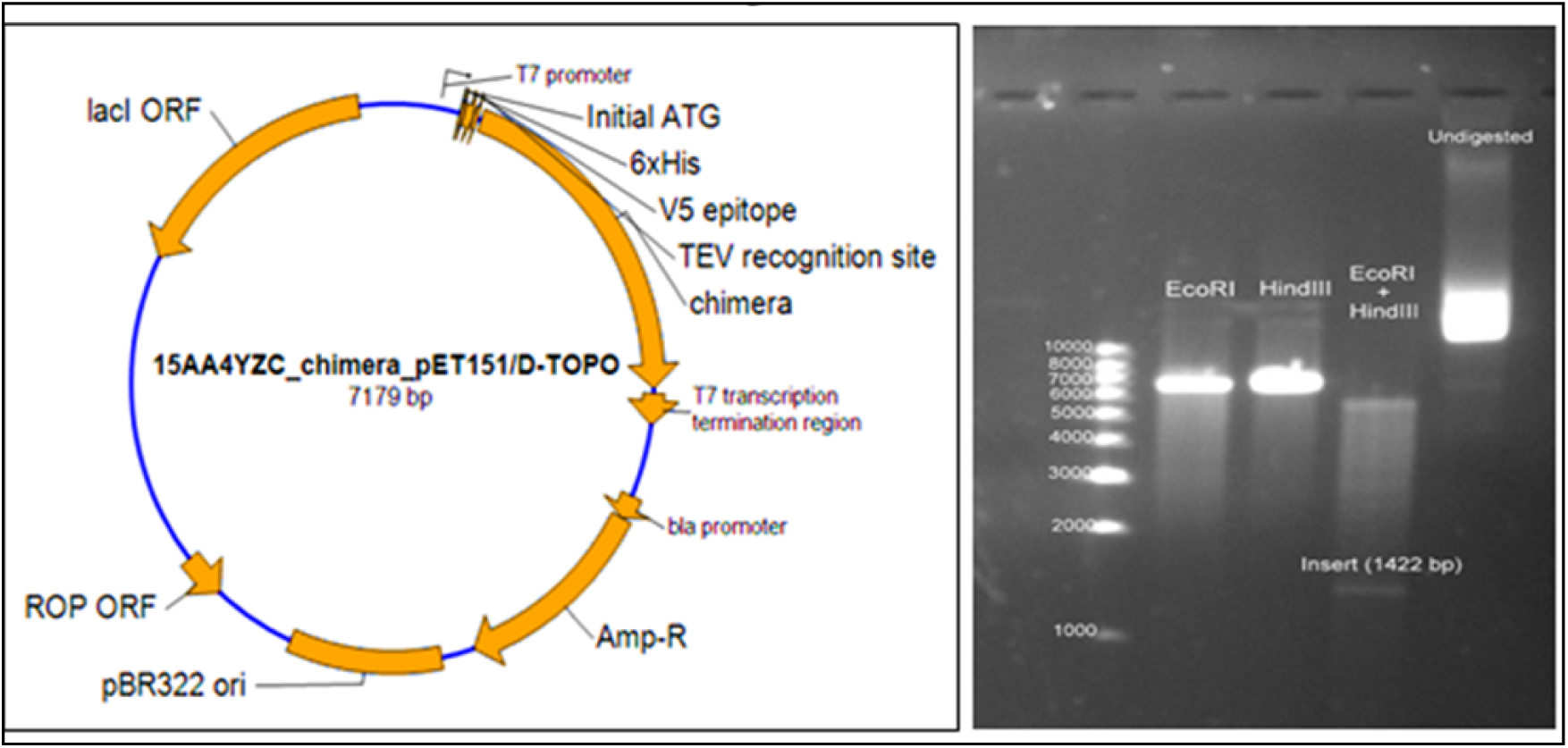
Plasmid map and restriction digestion.

**Figure 13:**
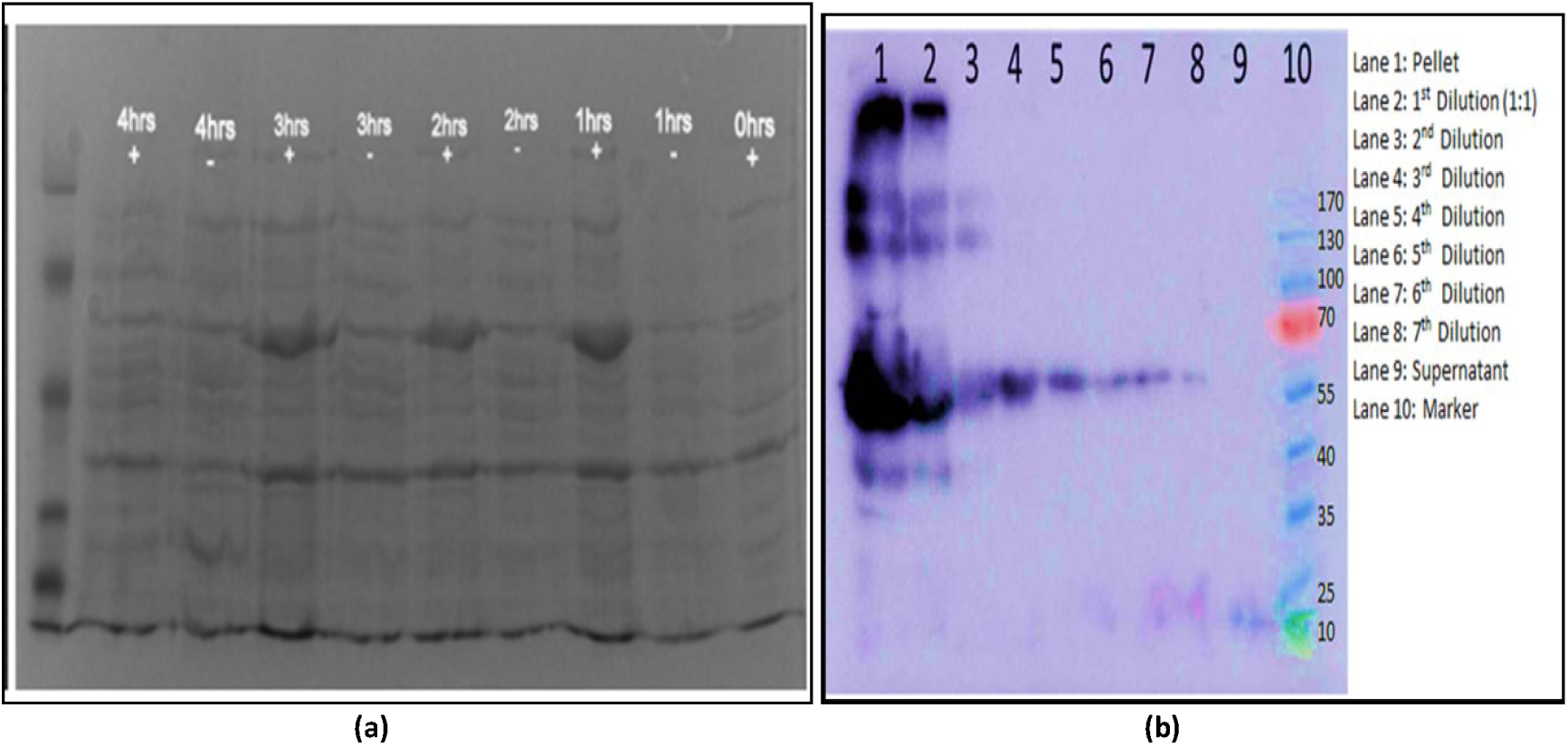
(a) Induction with 1mM IPTG from zero to 4 hours (SDS PAGE); (b) His probing of the His-tagged chimeric PKC.

### Negative autoregulatory circuit design and its quasipotential landscape

The negative autoregulatory circuit (**Figure 14 (a)**) *in silico* simulation was done in Berkeley Madonna (v 8.3), using a modified equation inspired from the toggle switch of Gardner et al. ^44^. The circuit has a single negative feedback loop in the form of LacI repressor protein, whose activity dictates the level of Chimeric PKC in the system. The synthetic circuit and the equation for the circuit are shown in **Figure 14 (b)**. The simulation shows the oscillations generated due to impulse of IPTG at 24 hour time point at a concentration of 1mM.

**Figure 14:**
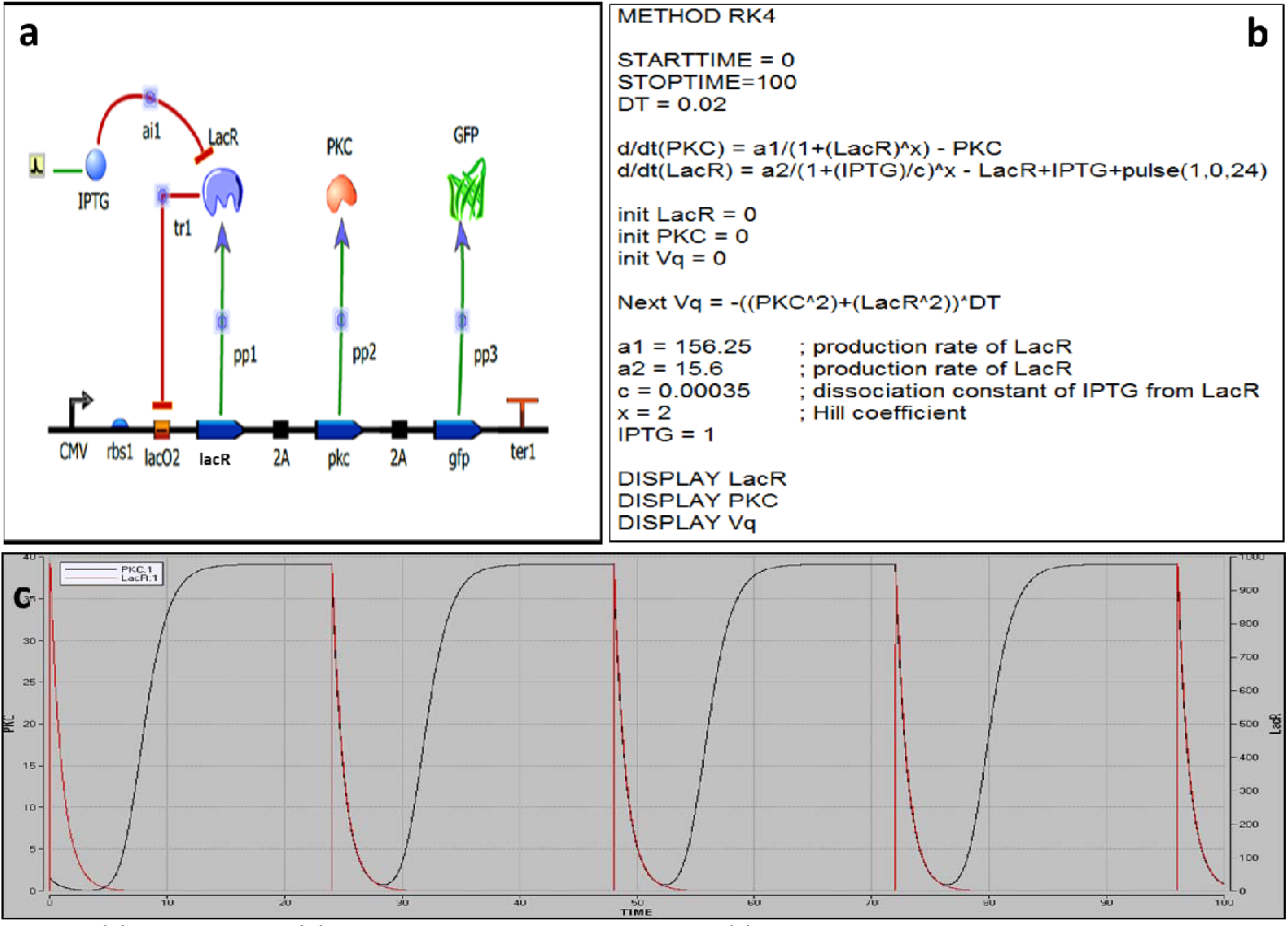
(a) Circuit Design, (b) equation in Berkeley Madonna; and (c) simulation results showing oscillations between the LacR and PKC due to the pulse of IPTG.

The time series data generated in Berkeley Madonna was further analyzed in R package, which shows that the system has two attractor states, i.e. either ON or OFF state (**Figure 15 (a)**). Attractor states suggest the stable gene expression pattern of the Boolean network. This system of a single negative autoregulator can oscillate between the two stable states ON and OFF in the presence or absence of the inducer IPTG, respectively. This helps in tuning the circuit response to the IPTG concentration. The network wiring (**Figure 15 (b)**) shows how the components are wired with respect to each other i.e. LacI R regulates itself and the expression of chimeric PKC. A state transition network depicts the binary representation of the attractors and the transitions between the attractor states. The total number of transition states in the circuit was 2, corresponding to the ON and OFF attractor states. Nullcline for the two ODEs of the synthetic circuit shows an asymptotically stable state i.e. the initial conditions of the system remain close to the initial condition (limiting behavior) and eventually converge to the equilibrium. For the synthetic circuit, the initial condition of the system is in the OFF state, which is turned ON with the impulse of the inducer IPTG. The system returns to the initial OFF state as the IPTG concentration falls from 1mM to zero nM and the system is ready for another impulse of IPTG. All the trajectory of the system converge to the center of the nullcline, at a single point indicating that the system can be in either the ON or OFF state and never in between i.e. intermediate state doesn’t exist. The point of intersection of nullclines is the saddle point or the equilibrium point.

**Figure 15:**
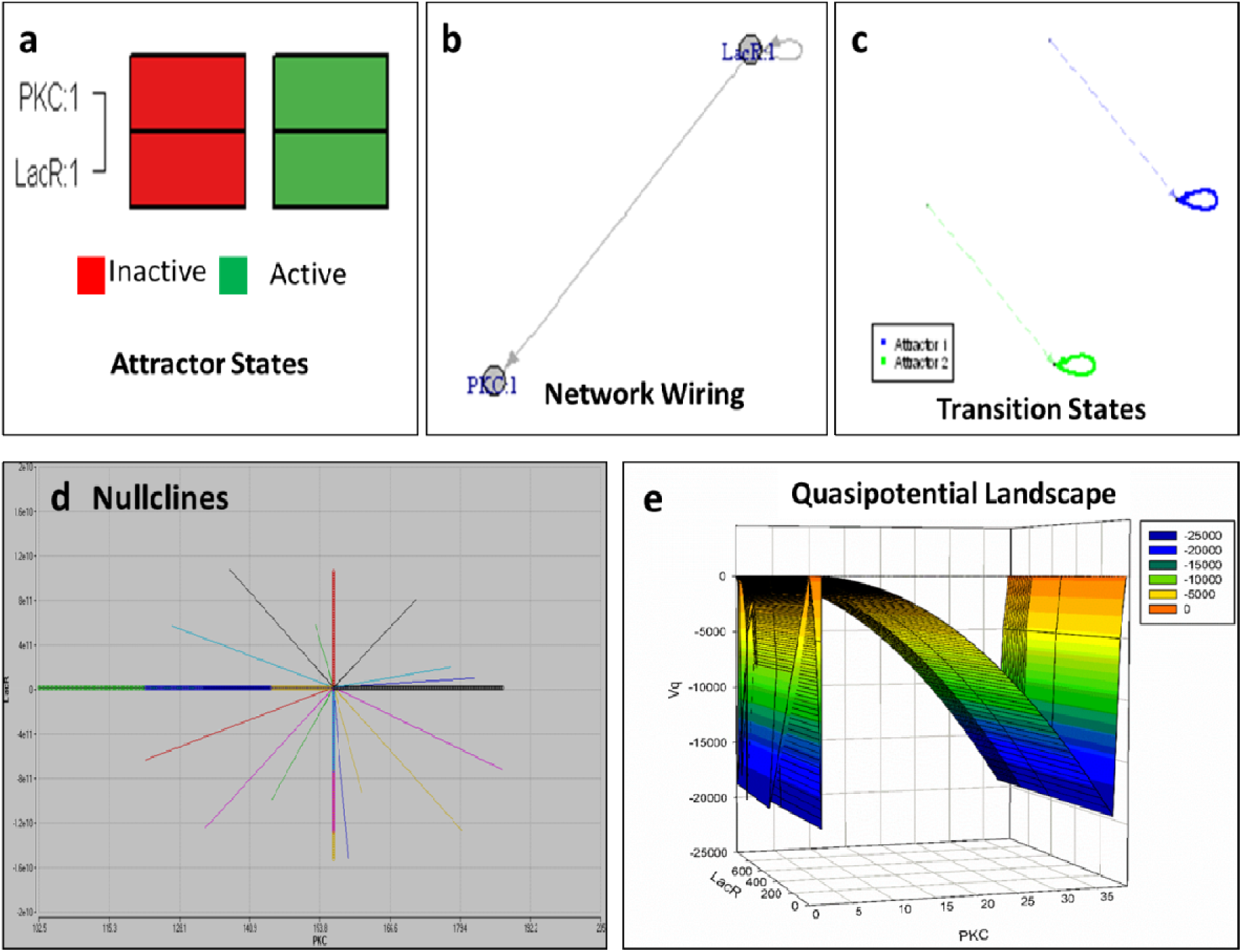
BoolNet Analysis.

The global dynamics of the system can be inferred by assigning potential to the attractor states. Hence the potential landscape states how forces on the system drive the system to the stable states. For the single negative auto regulatory circuit the force that drives the ON and OFF mechanism is the input of IPTG (inducer). The quasipotential landscape shows how the system crosses the thermodynamic barriers to move from one state to the other (**Figure 15 (e)**). Moreover, it can be visualized in **Figure 15 (e)** that the attractors remain in a state of low potential and upon addition of inducer, the system moves towards a higher concentration of attractors. Potential valleys correspond to the attractors and peaks to the transition states.

While running the GRENITS ^40^ package in R, gives us the convergence plots associated to the adequate convergence of the Markov Chain Monte Carlo (MCMC), which is crucial for further analysis. If the convergence has not been reached, the results are not trustworthy. For the synthetic circuit there is a perfect convergence of the plots for Gamma, B, Mu, Rho and Lambda which signifies the steady state dynamics and also the robustness of the designed synthetic circuit.

### Transfection and *in vitro* working of the synthetic circuit

The chimeric PKC was introduced into the negative autoregulatory circuit and transfected into peritoneal macrophages. The transfection was successful with an efficiency of over 60% (**Figure 16**). The working of the circuit was established with 1mM IPTG induction for 24 hours and fluorescence of GFP protein, when the IPTG was withdrawn the circuit was in the off state i.e. no GFP production and hence no chimeric PKC (**Figure 17**). The mathematical model built is useful to study the cytokine response to anti-inflammatory and pro-inflammatory activities and the resulting dynamical patterns. The developed model has allowed us to predict these changes with varying control parameters. We proceeded in a phenomenological manner and has utilized a functional approach rather than biophysical and biochemical detailing of the model to describe the role of feedback loop insertion in our mathematical model. The model is analyzed from the dynamical system point of view. The steady state values of the model and the null-cline obtained reveals a monotonically increasing function with the variable associated. The asymptotically stable attractor, the saddle and the unstable equilibrium points are illustrated in the phase plane portrait obtained. It is noteworthy to observe a biphasic output with a large initial peak followed by a sustained plateau with smaller amplitude in the quasipotential landscape represented in this paper. This is in agreement with the experimental results. Although changing the model parameters alters the quantitative analyses, the obtained qualitative responses are the same. Thus, it can be said that the absolute values of the obtained results are less significant and thus the simulation results are qualitatively discussed with the experimental observations. The simulation results actually helped us to analyze the cross-talk points identified in our CD14-TNF-EGFR signalling model system.

**Figure 16:**
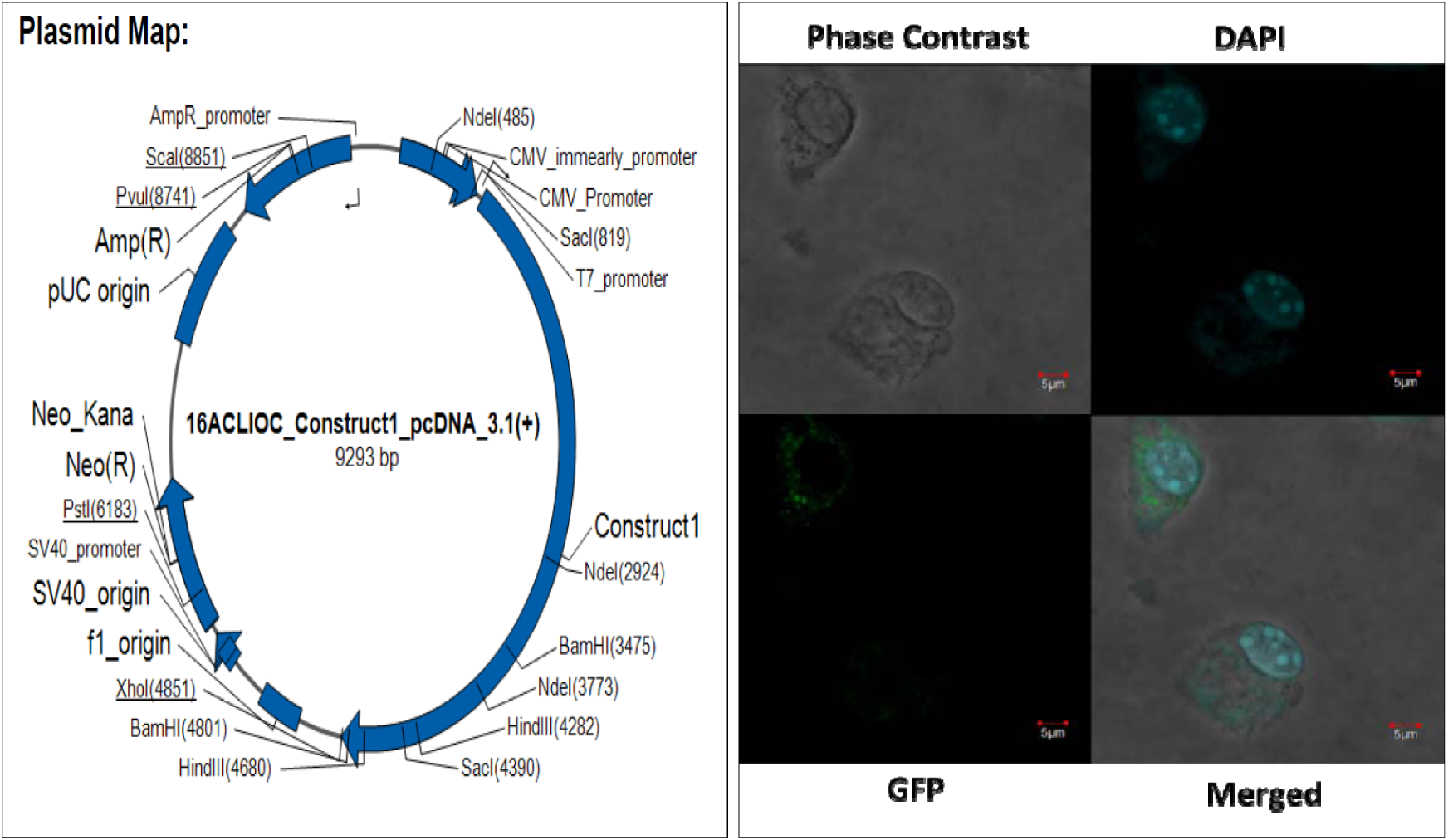
Synthetic construct design, transfection and expression of GFP.

**Figure 17:**
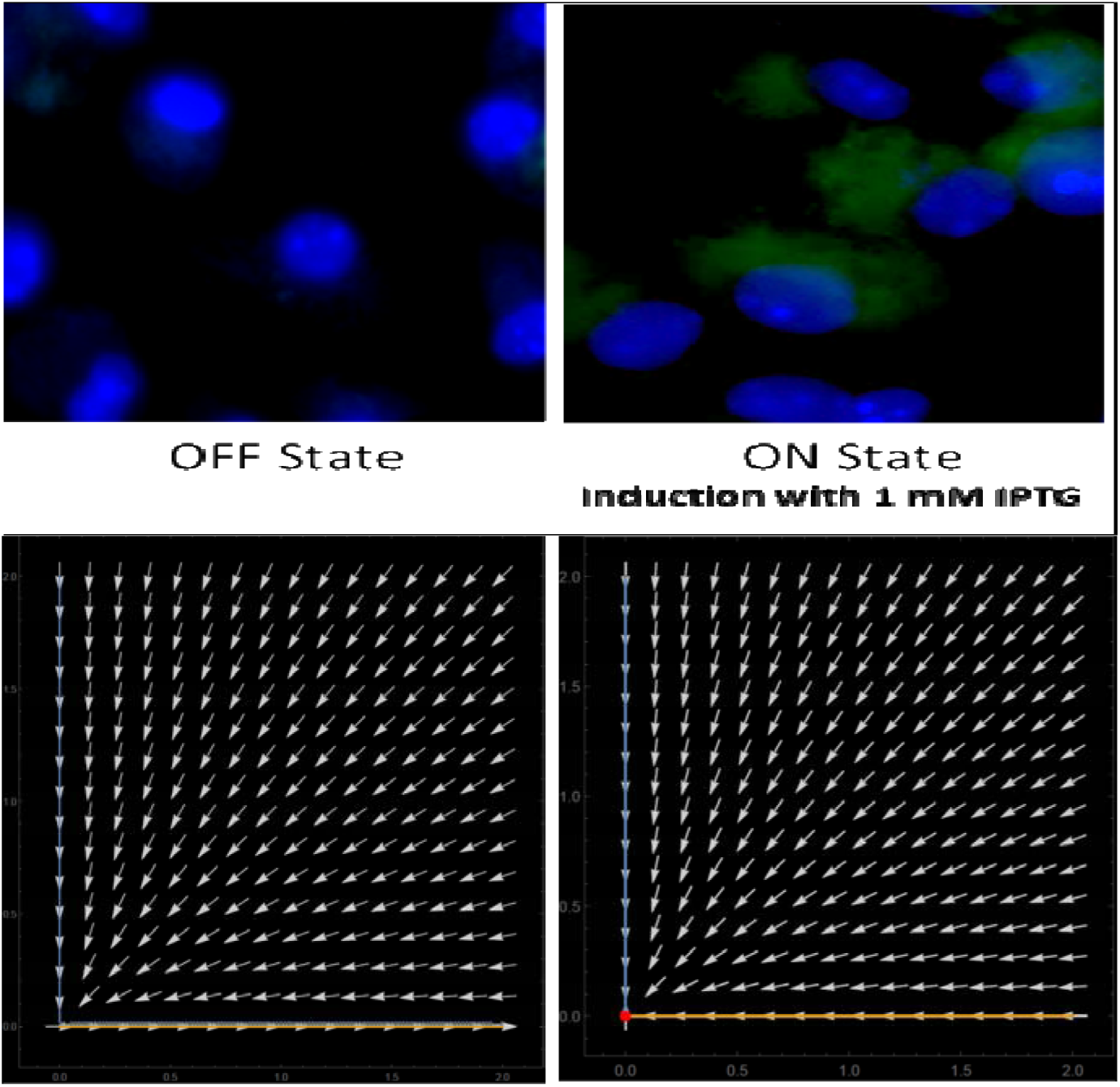
Working of the Circuit: Upper panel shows the OFF (no GFP expression) and ON (GFP expression in the presence of inducer IPTG) state; Lower plane shows the phase plane diagram for the same.

Western analysis shows thatthe chimeric PKC would be responsible for the phosphorylation of IKKb. As shown in **Figure 19 (a)**, the phosphorylated IKKb intensity is higher in the transfected induced lane compared to the transfected non-induced (mock) lane. This phenomenon was also observed in the confocal image (**Figure 19(b)**)

**Figure 19.**
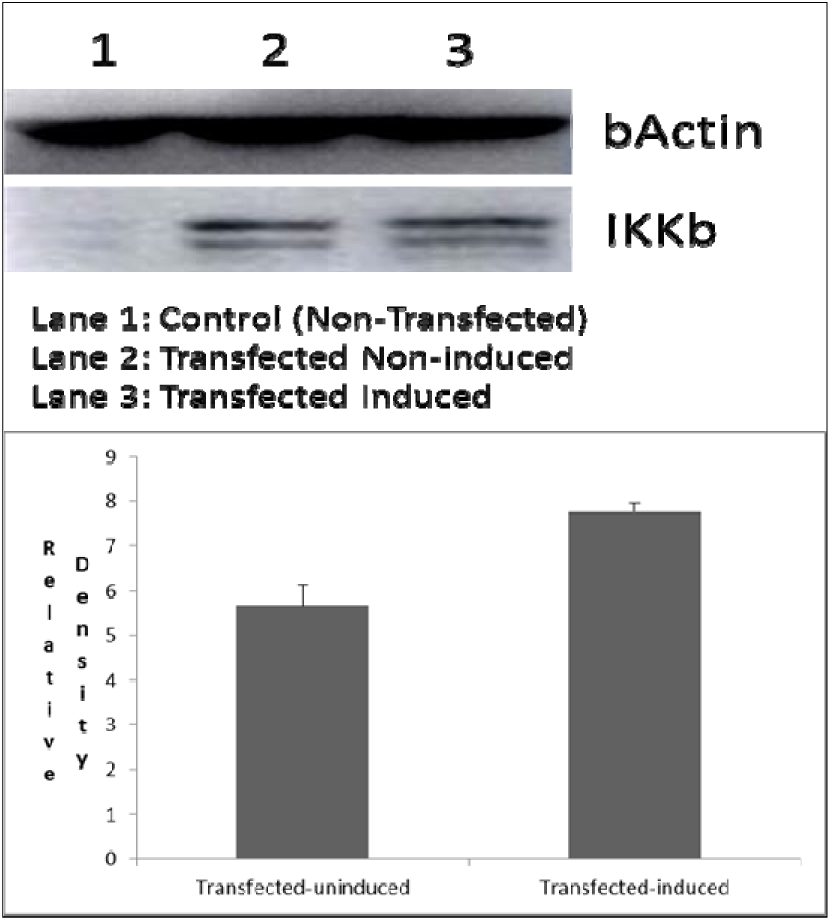
(a): Western blot showing the.

**Figure 19.**
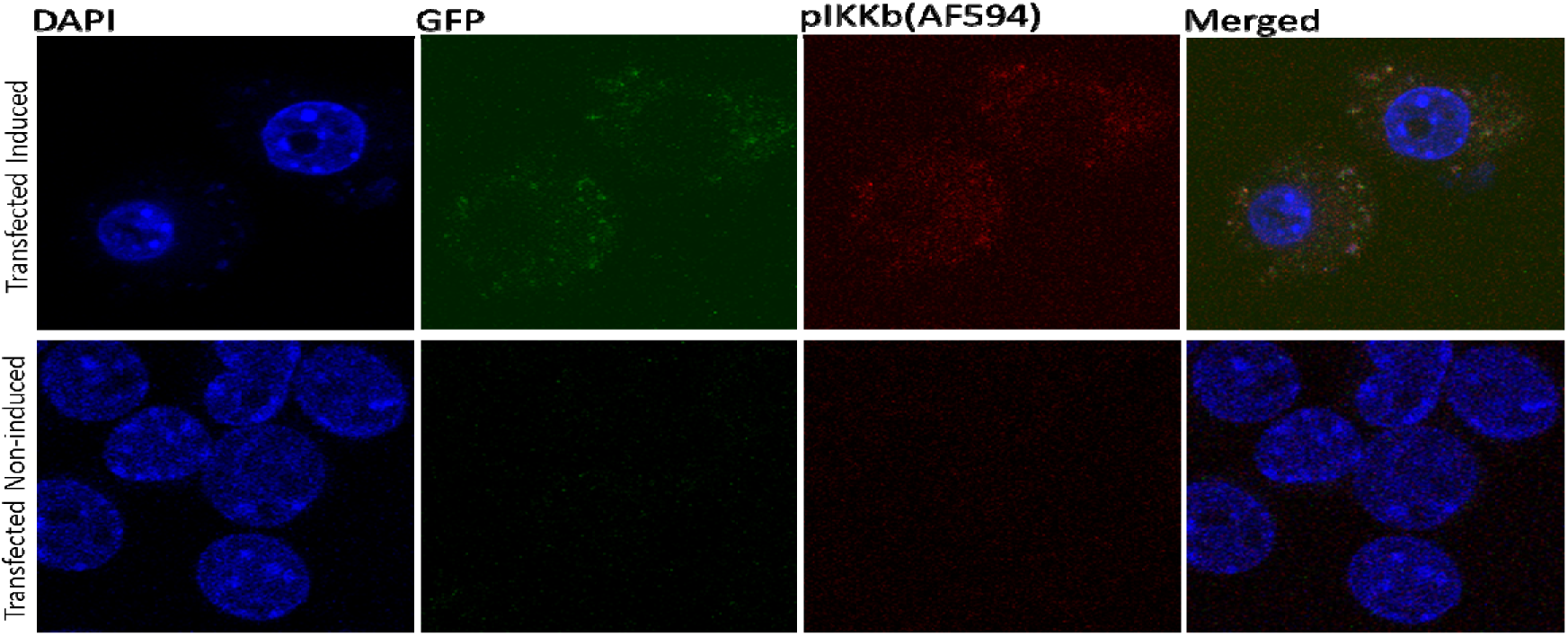
(b): Confocal Image of the transfected peritoneal macrophages showing the phosphorylation of Ikkb on induction with IPTG (1mM)

### Nitrite and Cytokine analysis

The percent infection and infectivity index of the *Leishmania* infected macrophages is shown in **Figure 20**. It is seen that in the transfected CTI and CTIM group, the percent infection stays close to 85% when compared to the I group with no significant change. While the infectivity index shows a reduced parasite load in the CTI and CTIM group when compared to the I group, suggesting that though there is decrease in the percent infection, there is reduction in the parasite load in the macrophages.

**Figure 20:**
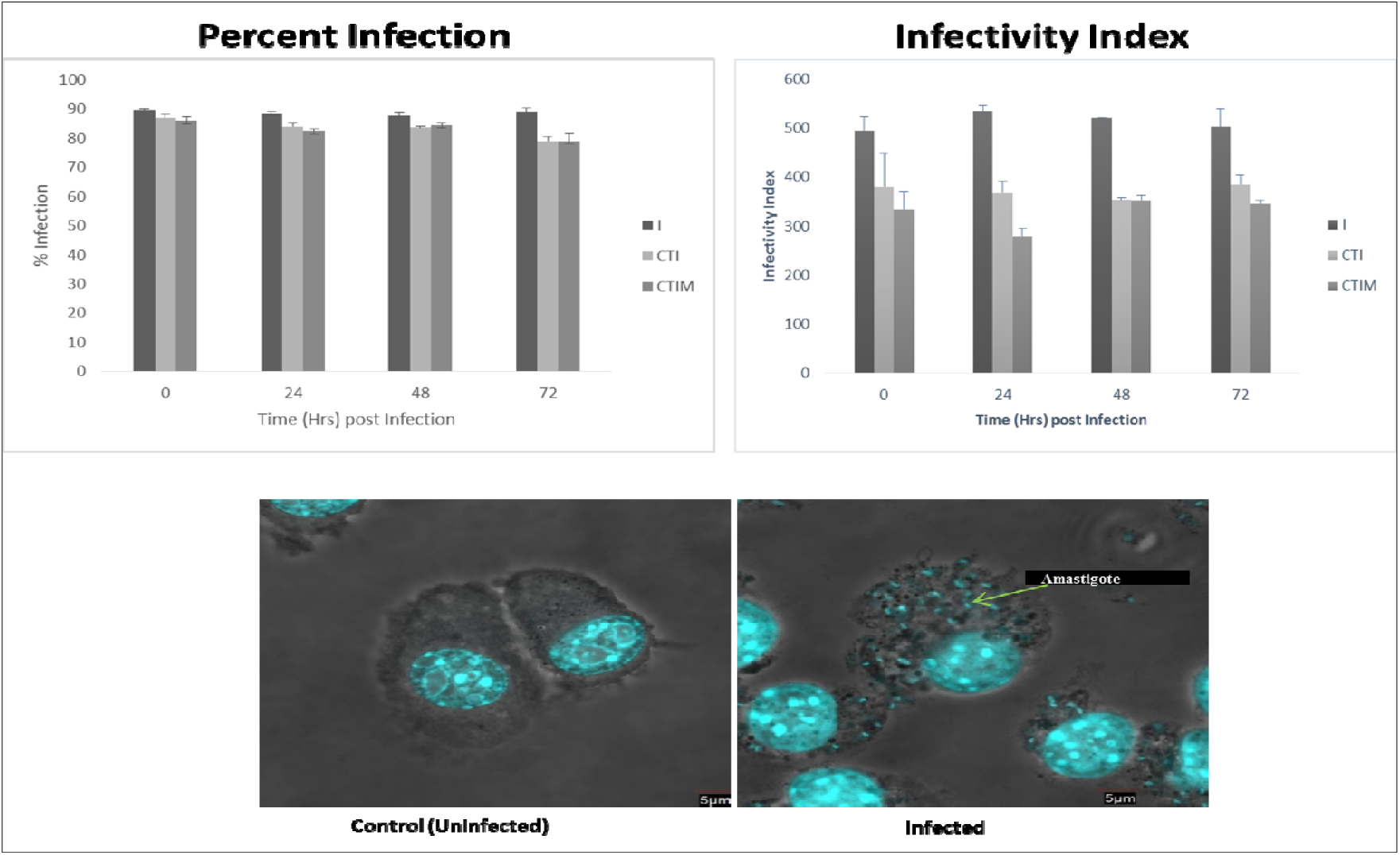
The percent infection and infectivity index of Leishmania infected macrophages.

The Nitrite estimation for the study groups is shown in **Figure 21.** As it can be observed, Infection does bring down the nitrite content which is elevated in the CT group. But in the CTI and CTMI group, there is decrease in the nitrite content, albeit not as low as in infection, suggesting that there is some NO production which is the cause for the lower parasite load in these groups.

**Figure 21:**
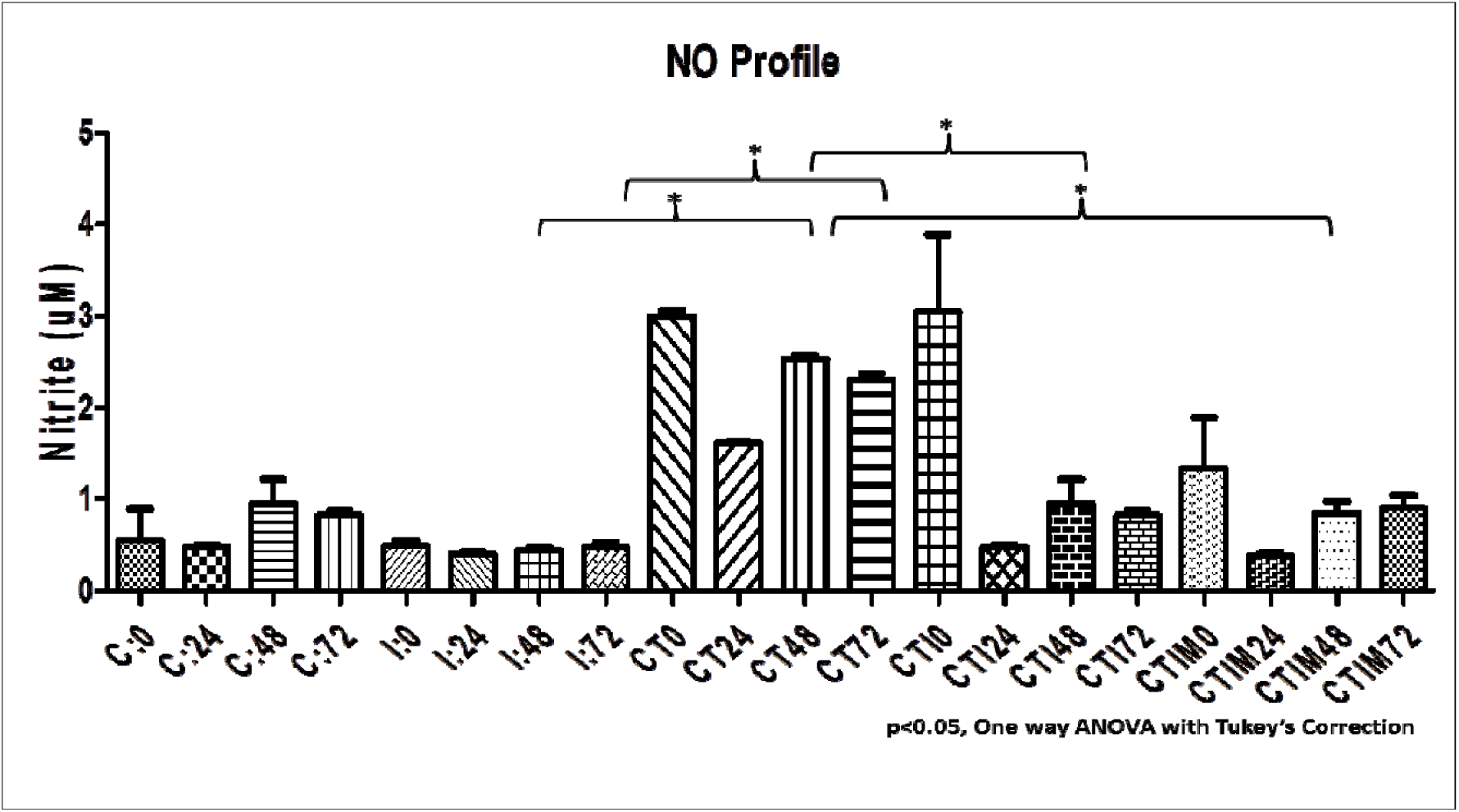
NO profile between the treatment groups.

While the cytokine analysis shows a non-significant change in the pro-inflammatory and anti-inflammatory cytokines (**Figure 22**). This could be due to the macrophage culture in isolation, which can be further modulated with T cell co-culture studies, for a disease resolving effect.

**Figure 22:**
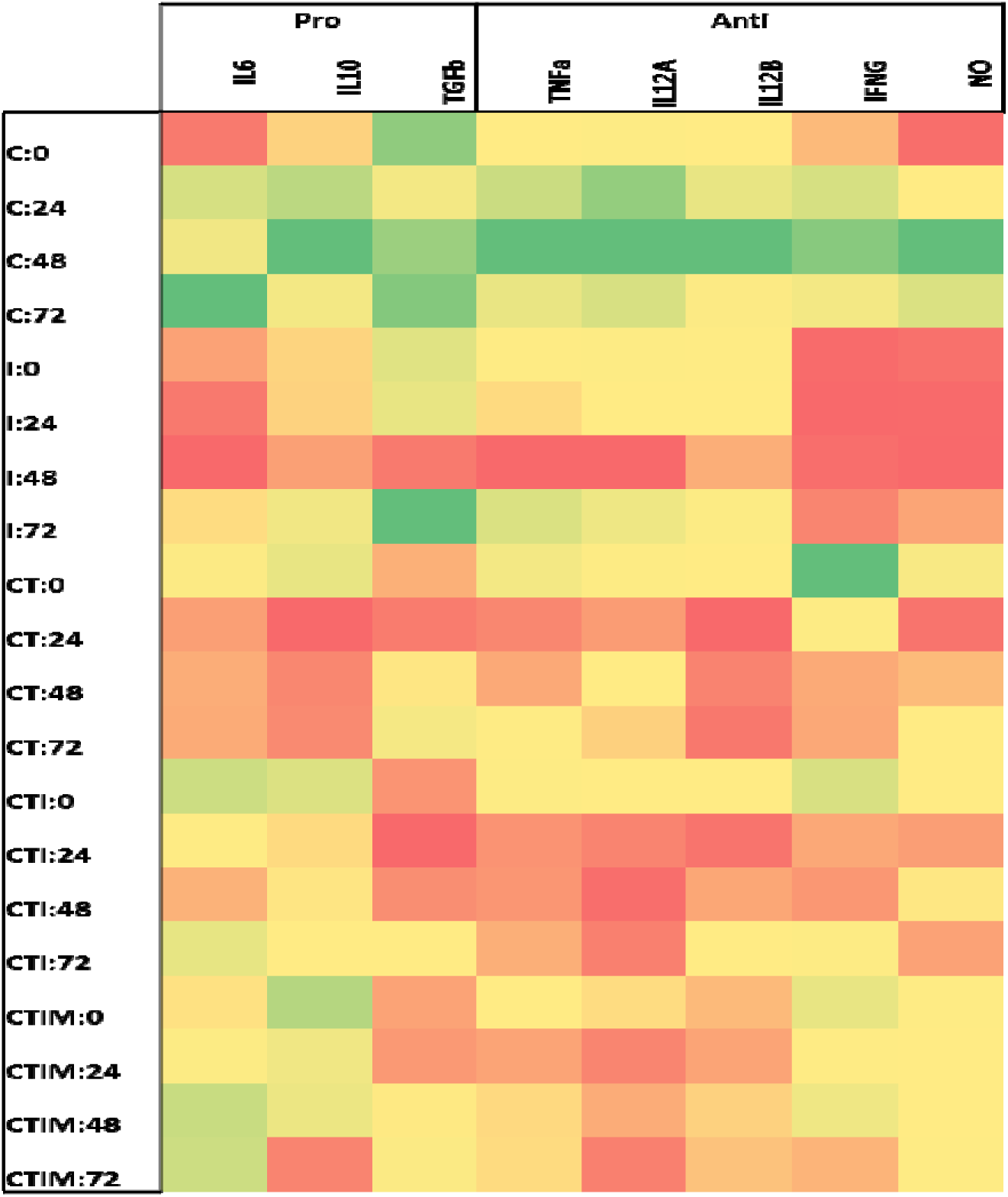
Cytokine analysis by qPCR.

## Discussion

Systems and synthetic biology have set their bench mark in the field of cancer and metabolic syndrome for therapeutics and diagnostics ^29–31,45^. There are no reports of synergistic application of systems and synthetic biology in the field of leishmaniasis. This work is the first of its kind in the leishmanial infection system. Though the system guided design of the synthetic circuit, shows a positive effect on the generation of pro inflammatory mediator NO, there is no significant effect on the synthesis of other pro inflammatory cytokines like TNFa, IL12 and IFNg. This system may be improved further for a desired modulatory effect by fine tuning the response of the synthetic circuit. Also, the system is only a macrophage system, may be a cell to cell communication is required to show a immune modulating effect. This could be achieved macrophage – T cell co culture studies.

Interfacing the computational and synthetic biology tools accelerate our understanding of complex biological systems and also our ability to quantitatively engineer cells that may aid as therapeutic device. Cells are basically engineered either to aid in various delivery platforms or to sense a specific molecule(s) that may distinguish the diseased vs. healthy state of the infection model system chosen for the study. The crux, here, lies into having a hierarchical architecture of the functional modules (parts assembled in a way to form an integrated biological system). The biochemical parameters related to transcription factor (TF) promoter binding is quantitatively determined and specifies which promoter drives the expression of which gene so that the network can be easily rewired. The system is robust to changes in parameter values as this is a context dependent system we have focused on, while laying the basic principle design of the synthetic signaling circuit.

The study began with a motivation when we identified that a negative auto regulatory module is embedded within a large transcriptional network. Like, electronic circuits are designed so that the connection of new inputs or outputs does not in any way impact the properties of a given module so are the synthetic biological circuits. The promoter binds and sequesters protein that has pleiotropic effects over the entire network. If the total promoter concentration is small compared to the TF-promoter equilibrium constant, sequestration effects are negligible and we do expect the total promoter concentration to be reliable. The promoter concentration ensures the synthetic device component reliability very much like the electronic circuits.

In addition, by designing synthetic systems which are translated and transcribed by completely orthogonal gene expression machinery offers another way of taking advantage of a host cell without interfering with its components. The effects a host plays on the synthetic circuit behaviour should be either minimized or where applicable should be quantified in order to characterize and validate parts/modules so as to minimize crosstalk between an engineered circuit and a host’s machinery. Thus, in the cellular context the designed synthetic circuit has better orthogonality (decoupling) with higher predictability and robustness.

In nutshell, we present paradigms for the design, construction and validation of robust synthetic circuits which may act as an implantable therapeutic device with more scale-ups or reuse of the modules in future.

## Acknowledgements

Milsee Mol acknowledges Senior Research fellowship from Council of Scientific and Industrial Research (CSIR), Ministry of Human Resource and Development, India. The authors thank Department of Biotechnology (DBT), Ministry of Science and Technology, for the funding. We take this opportunity to thank our Director, National Centre for Cell Science, Pune for supporting the Bioinformatics and High Performance Computing Facility (BHPCF) at NCCS, Pune, India. The support rendered by Proteomics Facility, NCCS and Confocal Microscopy Facility, NCCS are too acknowledged.

## References

1. Olliaro, P. L. & Bryceson, A. D. M. Practical progress and new drugs for changing patterns of leishmaniasis. Parasitol. today 9, 323–328 (1993).

2. Tiuman, T. S., Santos, A. O., Ueda-Nakamura, T., Dias Filho, B. P. & Nakamura, C. V. Recent advances in leishmaniasis treatment. Int. J. Infect. Dis. 15, e525--e532 (2011).

3. Organization, W. H. & others. Control of the leishmaniases: report of a meeting of the WHO Expert Commitee on the Control of Leishmaniases, Geneva, 22-26 March 2010. (2010).

4. Croft, S. L. & Olliaro, P. Leishmaniasis chemotherapy—challenges and opportunities. Clin. Microbiol. Infect. 17, 1478–1483 (2011).

5. Meheus, F. et al. Cost-effectiveness analysis of combination therapies for visceral leishmaniasis in the Indian subcontinent. PLoS Negl Trop Dis 4, e818 (2010).

6. Sindermann, H. et al. Miltefosine (Impavido): the first oral treatment against leishmaniasis. Med. Microbiol. Immunol. 193, 173–180 (2004).

7. Seifert, K. & Croft, S. L. In vitro and in vivo interactions between miltefosine and other antileishmanial drugs. Antimicrob. Agents Chemother. 50, 73–79 (2006).

8. Prajapati, V. K. et al. Targeted killing of Leishmania donovani in vivo and in vitro with amphotericin B attached to functionalized carbon nanotubes. J. Antimicrob. Chemother. dkr002 (2011).

9. Smith, A. C., Yardley, V., Rhodes, J. & Croft, S. L. Activity of the novel immunomodulatory compound tucaresol against experimental visceral leishmaniasis. Antimicrob. Agents Chemother. 44, 1494–1498 (2000).

10. Roychoudhury, J., Sinha, R. & Ali, N. Therapy with sodium stibogluconate in stearylamine-bearing liposomes confers cure against SSG-resistant Leishmania donovani in BALB/c mice. PLoS One 6, e17376 (2011).

11. Shio, M. T. et al. Drug delivery by tattooing to treat cutaneous leishmaniasis. Sci. Rep. 4, 4156 (2014).

12. Solomon, M. et al. Liposomal amphotericin B in comparison to sodium stibogluconate for cutaneous infection due to Leishmania braziliensis. J. Am. Acad. Dermatol. 56, 612–616 (2007).

13. Van Zandbergen, G., Hermann, N., Laufs, H., Solbach, W. & Laskay, T. Leishmania promastigotes release a granulocyte chemotactic factor and induce interleukin-8 release but inhibit gamma interferon-inducible protein 10 production by neutrophil granulocytes. Infect. Immun. 70, 4177–4184 (2002).

14. Nylen, S., Maasho, K., Söderström, K., Ilg, T. & Akuffo, H. Live Leishmania promastigotes can directly activate primary human natural killer cells to produce interferon-gamma. Clin. Exp. Immunol. 131, 457–467 (2003).

15. Laurenti, M. D. et al. The role of natural killer cells in the early period of infection in murine cutaneous leishmaniasis. Brazilian J. Med. Biol. Res. 32, 323–325 (1999).

16. Stenger, S., Thüring, H., Röllinghoff, M. & Bogdan, C. Tissue expression of inducible nitric oxide synthase is closely associated with resistance to Leishmania major. J. Exp. Med. 180, 783–793 (1994).

17. Nandan, D. & Reiner, N. E. Attenuation of gamma interferon-induced tyrosine phosphorylation in mononuclear phagocytes infected with Leishmania donovani: selective inhibition of signaling through Janus kinases and Stat1. Infect. Immun. 63, 4495–4500 (1995).

18. Scott, P. IFN-gamma modulates the early development of Th1 and Th2 responses in a murine model of cutaneous leishmaniasis. J. Immunol. 147, 3149–3155 (1991).

19. Kopf, M. et al. IL-4-deficient Balb/c mice resist infection with Leishmania major. J. Exp. Med. 184, 1127–1136 (1996).

20. Noben-Trauth, N., Lira, R., Nagase, H., Paul, W. E. & Sacks, D. L. The relative contribution of IL-4 receptor signaling and IL-10 to susceptibility to Leishmania major. J. Immunol. 170, 5152–5158 (2003).

21. Mylonas, K. J., Nair, M. G., Prieto-Lafuente, L., Paape, D. & Allen, J. E. Alternatively activated macrophages elicited by helminth infection can be reprogrammed to enable microbial killing. J. Immunol. 182, 3084–3094 (2009).

22. Arnold, L. et al. Inflammatory monocytes recruited after skeletal muscle injury switch into antiinflammatory macrophages to support myogenesis. J. Exp. Med. 204, 1057–1069 (2007).

23. Mounier, R. et al. AMPK$α$1 regulates macrophage skewing at the time of resolution of inflammation during skeletal muscle regeneration. Cell Metab. 18, 251–264 (2013).

24. Cohen, P. The development and therapeutic potential of protein kinase inhibitors. Curr. Opin. Chem. Biol. 3, 459–465 (1999).

25. Bagnyukova, T. et al. Chemotherapy and signaling: How can targeted therapies supercharge cytotoxic agents? Cancer Biol. Ther. 10, 839–853 (2010).

26. MacKenzie, S. H., Schipper, J. L. & Clark, A. C. The potential for caspases in drug discovery. Curr. Opin. Drug Discov. Devel. 13, 568–76 (2010).

27. Clark, K. et al. Phosphorylation of CRTC3 by the salt-inducible kinases controls the interconversion of classically activated and regulatory macrophages. Proc. Natl. Acad. Sci. 109, 16986–16991 (2012).

28. Kaminska, B. & Swiatek-Machado, K. Targeting signaling pathways with small molecules to treat autoimmune disorders. Expert Rev. Clin. Immunol. 4, 93–112 (2008).

29. Rössger, K., Charpin-El-Hamri, G. & Fussenegger, M. A closed-loop synthetic gene circuit for the treatment of diet-induced obesity in mice. Nat. Commun. 4, 2825 (2013).

30. Ye, H., Baba, M. D., Peng, R. & Fussenegger, M. Device Enhances Blood-Glucose Homeostasis in Mice.

31. Kemmer, C. et al. Self-sufficient control of urate homeostasis in mice by a synthetic circuit. Nat. Biotechnol. 28, 355–60 (2010).

32. Mol, M., Patole, M. S. & Singh, S. Immune signal transduction in leishmaniasis from natural to artificial systems: role of feedback loop insertion. Biochim. Biophys. Acta (BBA)-General Subj. 1840, 71–79 (2014).

33. Deb, K., Pratap, A., Agarwal, S. & Meyarivan, T. A fast and elitist multiobjective genetic algorithm: NSGA-II. IEEE Trans. Evol. Comput. 6, 182–197 (2002).

34. Eswar, N. et al. Comparative protein structure modeling using Modeller. Current protocols in bioinformatics **Chapter 5**, (2006).

35. Corpet, F. Multiple sequence alignment with hierarchical clustering. Nucleic Acids Res. 16, 10881–90 (1988).

36. Wiederstein, M. & Sippl, M. J. ProSA-web: Interactive web service for the recognition of errors in three-dimensional structures of proteins. Nucleic Acids Res. 35, 407–410 (2007).

37. Lovell, S. C. et al. Structure validation by C alpha geometry: phi,psi and C beta deviation. Proteins-Structure Funct. Genet. 50, 437–450 (2003).

38. Schrödinger, LLC. The {PyMOL} Molecular Graphics System, Version~1.8. (2015).

39. Bowers, K. et al. Scalable Algorithms for Molecular Dynamics Simulations on Commodity Clusters. ACM/IEEE SC 2006 Conf. 43–43 (2006). doi:10.1109/SC.2006.54

40. Morrissey, E. R. GRENITS : Gene Regulatory Network Inference Using Time Series Example: Network Inference For Simulated Data. 1–5 (2015).

41. Müssel, C., Hopfensitz, M. & Kestler, H. A. BoolNet— an R package for generation, reconstruction and analysis of Boolean networks. Bioinformatics 26, 1378–1380 (2010).

42. Bhattacharya, S., Zhang, Q. & Andersen, M. E. A deterministic map of Waddington’s epigenetic landscape for cell fate specification. BMC Syst. Biol. 5, 85 (2011).

43. Gasteiger, E. et al. Protein identification and analysis tools on the ExPASy server. (Springer, 2005).

44. Gardner, T. S., Cantor, C. R. & Collins, J. J. Construction of a genetic toggle switch in Escherichia coli. Nature 403, 339–342 (2000).

45. Lanitis, E. et al. Chimeric antigen receptor T cells with dissociated signaling domains exhibit focused antitumor activity with reduced potential for toxicity in vivo. Cancer Immunol. Res. 2013; 1: 43--53. doi: 10.1158/2326-6066.

